# Wound induced small-peptide mediated signalling cascade regulated by OsPSKR, dictates balance between growth and defense in rice

**DOI:** 10.1101/2023.06.20.545841

**Authors:** C.Y. Harshith, Avik Pal, Monoswi Chakraborty, Ashwin Nair, Steffi Raju, P. V. Shivaprasad

## Abstract

Wounding is a general stress in plants that results from various pest and pathogenic infections in addition to environment induced mechanical damages. Plants have sophisticated molecular mechanisms to recognize and respond to pests and pathogens. Although several molecules such as phytohormones, peptides and receptors have been attributed to wound responses in dicots, such mechanisms for monocots probably having distinct wound responses are less understood. Here, we show the involvement of two distinct categories of temporally separated, endogenously derived peptides, namely, plant elicitor peptides (PEPs) and phytosulfokine (PSK), that mediate wound responses in rice. These peptides trigger a dynamic signal relay in which a novel receptor kinase named OsPSKR played a major role. OsPSKR perceived PSK ligand, acting in association with a co-receptor OsSERK1, to activate downstream responses in a kinase activity-dependent manner. Perturbation of OsPSKR expression in rice led to compromised development and constitutive autoimmune phenotypes. These results suggested that OsPSKR maintains the trade-off between growth and exaggerated defense responses, both during homeostasis and wounding. Collectively, these findings indicate the presence of a stepwise peptide-mediated signal relay that regulates the transition from defense to growth upon wounding in monocots.

**One line summary:** Endogenous peptide signalling initiated wound responses through a receptor-like kinase OsPSKR to maintain the balance between growth and defense responses.

## Introduction

Plants encounter various pests, pathogens and mechanical damages in their natural ecosystems. These encounters result in the loss of cell wall integrity (CWI). Cell wall damages are constantly monitored by various CWI sensors and any deviation leads to various molecular and metabolic responses (Engelsdorf *et al*., 2018). Persistent or unattended wound sites serve as entry points for various secondary infections and can affect the fitness of plants. Therefore, plants have evolved tightly coordinated molecular signalling networks to sense and respond to wounding. The earliest molecular signalling activated by wounding is part of general stress responses that are very similar to pathogen-derived pattern triggered immune (PTI) responses. In the event of wounding, plants activate multitude of molecular responses to initiate defense signalling that include release of damage associated molecular patterns (DAMPs) from the damaged cells, activation of Ca_2+_ based signalling cascade, accumulation of reactive oxygen species (ROS) in the adjacent cells and activation of systemic signalling mediated by various phytohormones (Savatin *et al*., 2014; Vega-Muñoz *et al*., 2020). These responses eventually culminate in active blockage of wound site and promotion of further growth/regeneration (Hernández-Coronado *et al*., 2022). Although most of the early wound responses are similar between monocots and dicots, monocot wound responses are not understood in detail.

Dicots have remarkable regeneration abilities including *de novo* organogenesis (Hu *et al*., 2017; Zhang *et al*., 2019; Liang *et al*., 2023). Wounding triggers *de novo* root formation in *Arabidopsis* by coordinated action of jasmonic acid (JA) and auxin phytohormones (Zhang *et al*., 2019). Auxin mediated signalling also activates vascular regeneration post wounding in *Arabidopsis* that helps in wound healing (Radhakrishnan *et al*., 2020). However, monocots lack wound-induced regeneration ability and wound repair process is distinct in comparison to that of dicots (Liang *et al*., 2023). Upon injury, monocot leaf blades display minimal cell death and initiate growth, probably owing to cell wall remodelling and cell elongation. However, the molecular mechanism involved in wound responses among monocots that have been subjected to herbivory over the course of evolution is not well understood.

Major wounding events are a direct result of insect herbivory. Plants deter insect herbivores through various molecular responses and these responses are largely overlapping between wounding and herbivory (Erb & Reymond, 2019). Insect herbivory induced responses have been thought to be activated by insect-derived cues referred to as herbivore associated molecular patterns (HAMPs). However, so far, only one instance of HAMP-induced defense activation has been reported from cowpea where inceptin was found to trigger defense responses through a receptor like protein (RLP) called INCEPTIN RECEPTOR (Steinbrenner *et al*., 2020).

Signals emanating from endogenous sources play a major role in the activation of various signalling pathways across organisms. Plants code for a plethora of non-functional precursor proteins which when induced and processed, give rise to bioactive peptides, that are further capable of activating receptor mediated signalling pathways (Tavormina *et al*., 2015; Li *et al*., 2020). Molecular patterns derived from endogenous source are referred to as DAMPs, and these patterns activate their cognate receptors, eventually inducing innate immunity (Tanaka & Heil, 2021). Since effects of mechanical wounding and insect herbivory have been studied together, many DAMPs have been identified as major immune triggers for both the responses (Pearce *et al*., 1991; Huffaker *et al*., 2006). Among these, systemin is an 18 amino acid (aa) long peptide hormone specific to *Solanaceae* family. Systemin is derived from its non-functional precursor named prosystemin and it was among the first identified DAMPs in response to wounding in dicots (Pearce *et al*., 1991; Ryan & Pearce, 1998). Other DAMPs include plant elicitor peptides (PEPs) that are well-conserved across plant species and have been shown to be activated upon several cues including wounding (Huffaker *et al*., 2006, 2011, 2013; Trivilin *et al*., 2014; Lori *et al*., 2015; Shinya *et al*., 2018; Poretsky *et al*., 2020).

PEPs are 23 aa long with varying sequences across different families (Bartels *et al*., 2013; Lori *et al*., 2015). The transcriptional dynamics that leads to activation of downstream responses post PEP-perception are not well understood, apart from a study in which transcriptome analysis using multiple immune triggering peptides including AtPep1 was carried out (Bjornson *et al*., 2021). Each PEP is unique by sequence and they have distinct localization pattern. This prompts a thorough, species-specific as well as context-specific investigations of signalling regulated by PEPs.

Multiple studies have suggested the involvement of signal transitioning mediated by phytohormones such as JA and auxin (Zhang *et al*., 2019) in growth responses post-wounding. Peptide hormones also seem to play a key role in growth and development (Motomitsu *et al*., 2015), but their specific contributions to wound responses are relatively less understood. Among the peptide hormones, Phytosulfokine (PSK) is a sulphated pentapeptide hormone derived from secreted precursors and has been attributed in a multitude of roles including cell proliferation, tracheary element differentiation, vegetative growth, lateral root development and cell expansion (Matsubayashi & Sakagami, 1996; Matsubayashi *et al*., 1999, 2006; Sauter, 2015). PSK seems to be responding to wounding in *Arabidopsis* as PSK precursors were induced upon wounding (Matsubayashi *et al*., 2006; Loivamäki *et al*., 2010). Whether PSK signalling actively regulates downstream wound responses is unknown. PSK is recognized by leucine rich repeat receptor like kinases (LRR-RLK) across plants and direct evidence of the binding has been provided in *Arabidopsis*, carrot and tomato (Matsubayashi *et al*., 2002; Wang *et al*., 2015; Hohmann *et al*., 2017; Zhang *et al*., 2018). However, signalling regulated by PSK receptors is not thoroughly understood in monocots such as rice. Though there are 15 predicted potential PSK receptors (PSKRs) in rice (Nagar *et al*., 2020), the nature of signals they respond to, the ligands they bind to, or their downstream functions are largely unknown. Among these, OsPSKR1 has been attributed to disease resistance (Yang *et al*., 2019), while OsPSKR15 played a role in drought response (Nagar *et al*., 2022).

In this study, we find the role of two distinct categories of small peptides in wound perception and response in rice using time-course transcriptome analysis. We identified an early activation of *OsPROPEP3* upon wounding which is the precursor for OsPep3. Further, wounding and PEP-triggered signalling activated a DAMP, PSK. Using biochemical and *in silico* methods, we identified OsPSKR as a receptor for PSK. Using mutational studies, we found that OsPSKR plays an essential role in development since its mutants displayed very drastic phenotypes including sterility. We also identified its role in maintaining balance between growth and defense responses under homeostasis. During wound responses, OsPSKR assisted in transitioning from early defense responses to late growth/repair responses. These findings suggest the presence of a step-wise transcriptional program that relays wound-induced signals through a receptor-kinase that decides growth vs. defense responses in rice.

## Results

### Time course transcriptome screen identifies rapid induction of precursors of small peptides upon wounding

Among dicots, earliest wound responses include changes in membrane dynamics, calcium signalling, expression of peptide precursors, and rapid expression of various wound responsive genes (Savatin *et al*., 2014; Vega-Muñoz *et al*., 2020). A detailed, temporal progression of specific events among monocots upon wounding is lacking (Hu *et al*., 2017; Liang *et al*., 2023). To address this, a detailed multi-time point transcriptome analysis was carried out using six-week-old rice seedlings. Wounding was carried out by punching leaf blades at the edges at 2 cm intervals without affecting the midrib (Venu *et al*., 2010) and samples were collected at 15 min, 30 min, 2 h and 12 h post-wounding with appropriate unwounded controls (Fig. 1a).

**Figure 1:**
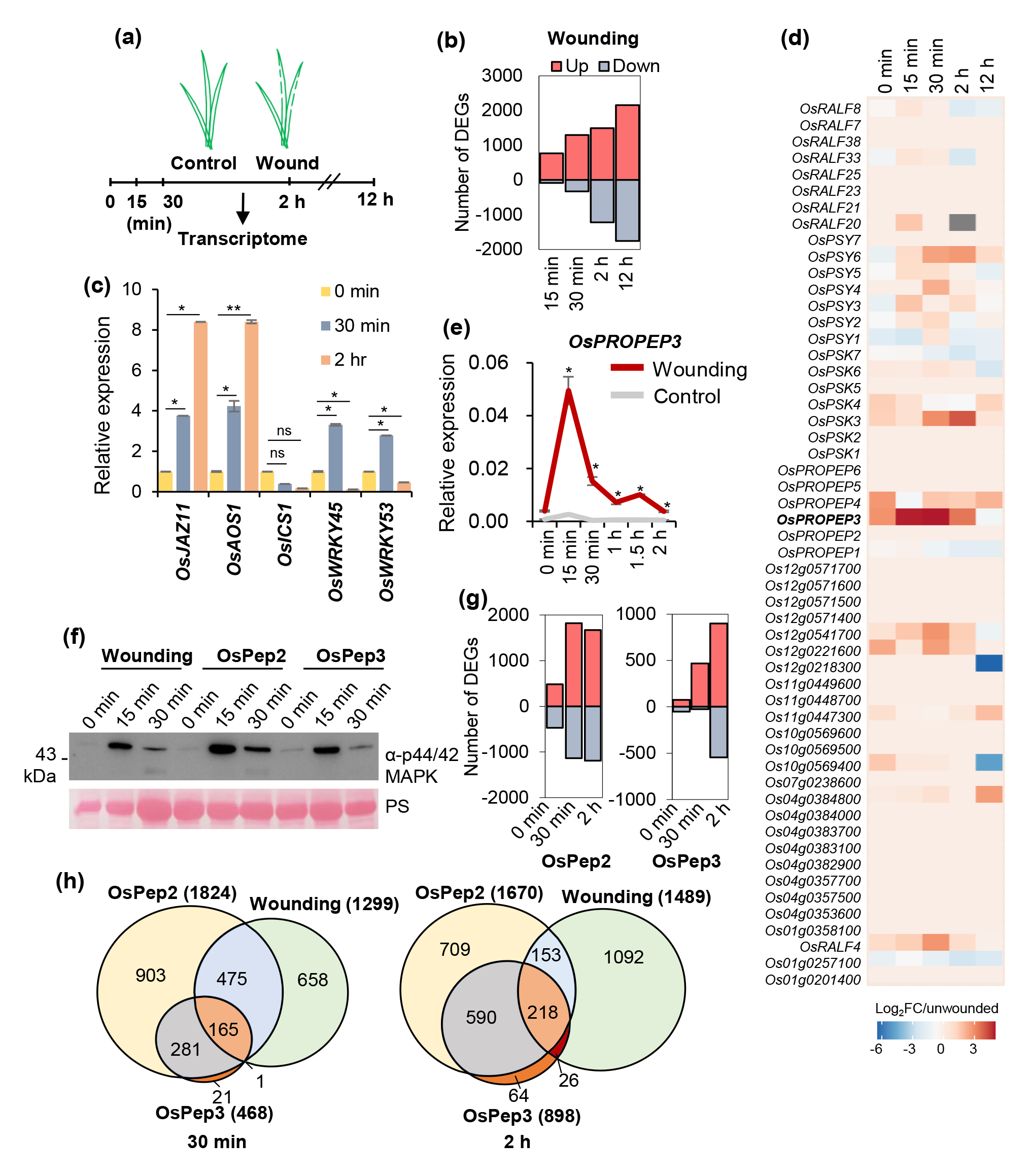
Time course transcriptome screen identifies rapid induction of precursors of small peptides upon wounding. **(a)** Scheme showing experimental time points. Six-week-old rice plants were mechanically injured, leaves collected at the indicated time points and subjected to transcriptome analysis. **(b)** Differential expression of genes (Log_2_FC≥1.5 for upregulated & Log_2_FC≤-1.5 for downregulated, p<0.05) upon wounding across time points. **(c)** RT-qPCR analysis of expression of *OsJAZ11*, *OsAOS1*, *OsICS1*, *OsWRKY45* and *OsWRKY53* relative to *OsActin* upon wounding, n=3, pairwise Student’s-*t* test with respect to 0 min samples, *: p<0.05, **: p<0.005. **(d)** Transcriptional changes of precursors of various DAMPs represented as Log_2_FC with respect to corresponding unwounded samples. **(e)** RT-qPCR analysis of expression of *OsPROPEP3* relative to *OsActin* upon wounding, n =3, pairwise Student’s-*t* test with respect to control samples, *: p<0.05. **(f)** MPK activation assay upon wounding, OsPep2 and OsPep3 treatments. **(g)** Number of significantly differentially regulated genes upon OsPep2 and OsPep3 treatments, Log_2_FC≥1.5 for upregulated & Log_2_FC≤-1.5 for downregulated, p<0.05. **(h)** Representation of the number of overlapping upregulated genes between wounding, OsPep2 and OsPep3 treatments, Log_2_FC ≥1.5, p<0.05. DEGs = Differentially expressed genes.

RNA-seq datasets were generated in biological duplicates and processed as described (Hari Sundar G *et al*., 2023). The number of DEGs increased over the time course after wounding indicating a gradually intensifying response to wounding (Fig.1b, Supplemental dataset 1). Studies in *Arabidopsis* have identified JA related genes as one of the earliest responding set of genes to wounding (Zhang *et al*., 2019). Similarly, wounding in rice also triggered JA signalling related genes at early time points (30 min and 2 h post wounding) in addition to other genes involved in signalling (Fig. S1a). JA pathway genes *OsJAZ11* and *OsAOS1* showed a very rapid response to wounding with increasing expression over time (Fig. 1c). However, in contrast to JA pathway genes, salicylic acid (SA) biosynthesis gene *OsICS1* did not respond to wounding as these two phytohormone pathways have been proposed to be antagonistic to each other (Nomoto *et al*., 2021). Two of the well-studied general stress responsive WRKY transcription factors- *OsWRKY45* and *OsWRKY53* also showed transcriptional upregulation upon wounding (Fig. 1c).

Many studies in diverse plants have shown the induction of small peptide coding genes either upon mechanical wounding or upon insect herbivory (Vega-Muñoz *et al*., 2020). We explored the transcriptional response of precursors of small peptide coding genes and their homologs (Li *et al*., 2020). These included a total of 54 small peptide coding precursors including PSKs, RALFs, PROPEPs and PSYs (Sauter, 2015; Sharma *et al*., 2016; Shinya *et al*., 2018; Tost *et al*., 2021). Among these precursors, two different peptide precursors that belong to distinct categories were found responding to wounding at different time points (Fig.1d, Supplemental dataset 2). *OsPROPEP3*, a gene that codes for a DAMP named OsPep3 showed a remarkable induction at a very early time point post wounding. *OsPROPEP3* was previously identified to be wound-responsive as well as responsive to herbivore derived oral secretion treatment in rice (Shinya *et al*., 2018). The transcriptional induction of *OsPROPEP3* was confirmed using RT-qPCR which showed a rapid increase in expression at 15 min and subsequent reduction beyond 30 min post wounding (Fig. 1e). Including OsPROPEP3, rice has a total of six predicted PROPEPs with varying sequences and give rise to bioactive peptides of 23 aa in length (Fig. S1b).

In order to understand if bioactive peptide derived from OsPROPEP3 can elicit wound responses, rice leaves were treated with OsPep3 peptide. Also, OsPep2 has been previously attributed to herbivore responses (Huffaker *et al*., 2013). Therefore, OsPep2 was also considered in subsequent experiments. MPK activation is one of the primary hallmarks of PTI activation and many studies have shown that PEPs can activate MPK signalling (Shinya *et al*., 2018; Zheng *et al*., 2018). In agreement with this, we observed a rapid activation of MPK post wounding as well as upon OsPep3 and OsPep2 treatments suggesting the activation of conventional PTI signalling (Fig. 1f). We also confirmed transcript upregulation of OsMPK3 upon wounding as well as OsPep3 and OsPep2 treatments (Fig. S1c & d). In order to identify the molecular responses activated by OsPep3 & OsPep2, and to assess the similarities and overlap between the responses activated by wounding and PEP treatment, we performed a time-point based transcriptome analysis after treating rice leaves using OsPep3 and OsPep2. Both OsPep3 and OsPep2 treatments led to differential expression of several hundred genes (Fig. 1g, Supplemental dataset 3).

In order to further understand if the genes responsive to OsPep3 and OsPep2 treatments are under wound response module, we overlapped genes responding to individual treatments with each other (Fig. 1h). We noticed that over 90% of the genes responsive to OsPep3 were also responsive to OsPep2, indicating the remarkable similarities between downstream responses upon PEP perception. Also, we observed a strong overlap of 49.2% (30 min) and 24.91% (2 h) between PEP responsive (upregulated) and wound responsive (upregulated) genes across two different time points, indicating that PEPs can trigger responses similar to wounding. It is important to note that the categories of genes responding to wounding changed over time, with early set of genes predominantly involved in JA signalling and late set of genes involved in ribosome subunit biogenesis and rRNA processing (Fig. S1a).

Both OsPep3 and OsPep2 treatments triggered similar set of genes that were upregulated upon 30 minutes post wounding (Fig. S1e). These results suggest that mechanical wounding induced a rapid transcriptional signalling including the induction of peptide coding genes. Bioactive peptides which are derivatives of precursor genes induced similar responses to that of wounding, indicating a surprising functional resemblance of downstream signalling.

### Wounding and wound-derived OsPep3 treatments activated PSK signalling

Previous studies have shown that transcriptional dynamics upon wounding involved transitioning from early defense responses to late growth responses (Zhang *et al*., 2019; Hernández-Coronado *et al*., 2022). To explore this possibility, we compared transcriptomes between early vs late time points. We observed a delayed upregulation of a precursor of a distinct category of small peptides, PSK (Fig. 2a, Supplemental dataset 4). There are seven predicted PSK coding genes in rice, and among which *OsPSK3* showed a remarkable induction at 2 h post-wounding. This response was also observed with OsPep2 and OsPep3 treatments, indicating PSK activation at a delayed time point might be under the regulation of a signal relay (Fig. 2a). RT-qPCR analysis of expression dynamics of *OsPSK3* upon wounding showed an induction 30 min post-wounding, indicating a possible peptide mediated temporal signalling relay either hierarchically or parallelly since *OsPROPEP3* induction was observed at 15 min post-wounding (Fig 2b, 1d). Induction of PSK upon wounding was also observed in *Arabidopsis* (Matsubayashi *et al*., 2006; Loivamäki *et al*., 2010), where it was perceived by an LRR-RLK named AtPSKR1 (Wang *et al*., 2015). The exact role and mechanism of PSK signalling in wound responses has not been explored. Induction of *OsPSK3* upon wounding as well as with OsPep3 treatment indicates specific activation of PSK mediated signalling. It is possible that PSK receptor is involved in wound responses in rice.

**Figure 2:**
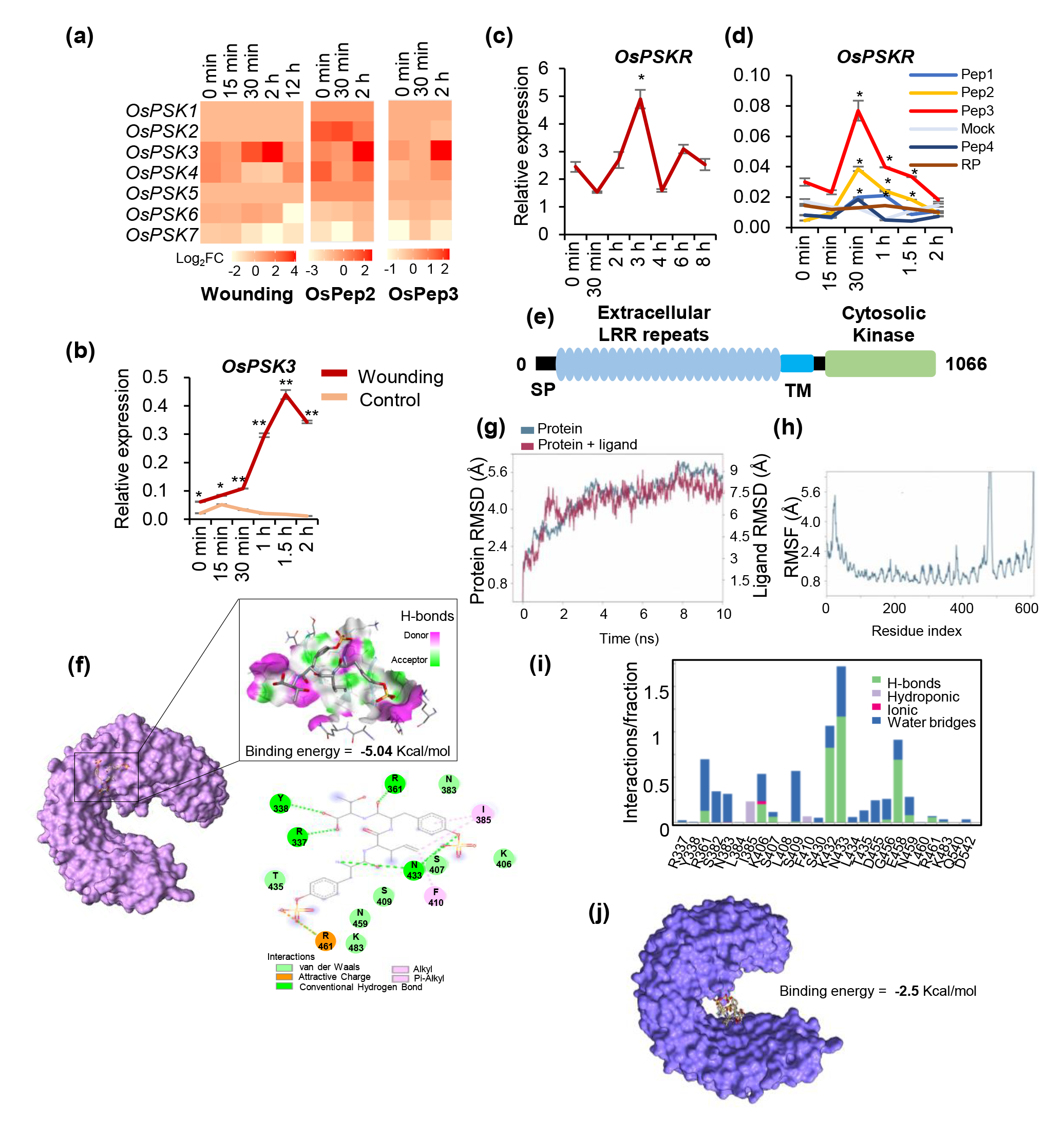
Wounding and wound-derived OsPep3 treatments activated PSK signalling. **(a)** Expression profile of predicted OsPSK precursors upon wounding, OsPep2 and OsPep3 treatment, represented as Log_2_FC with respect to corresponding unwounded samples. **(b)** RT-qPCR showing the transcriptional upregulation of *OsPSK3* relative to *OsActin* upon wounding, n=3. **(c)** RT-qPCR analysis of expression of *OsPSKR* relative to *OsActin* upon wounding across time points, n=9. **(d)** RT-qPCR analysis of expression of *OsPSKR* relative to *OsActin* upon peptide treatment across time points, n=3. **(e)** Domain architecture of OsPSKR. **(f)** Representation of ligand-receptor interaction obtained by docking. **(g,h)** RMSD, RMSF plots showing protein-ligand interactions in MD simulation. **(i)** MD simulation studies showing predicted residues taking part in ligand-receptor interaction. **(j)** Homology model of mutant OsPSKR (K432A & N433A) with unbound ligand. Statistics in **b,c,d**: pairwise Student’s-*t* test, *: p<0.05, **: p<0.005.

In order to ascertain probable role of a PSK receptor in transduction of signals upon wounding, we considered all 15 rice PSK receptors predicted previously (Yang *et al*., 2019; Nagar *et al*., 2020). Transcriptome analysis showed differential regulation of several predicted PSK receptor genes upon wounding as well as with OsPep3 treatment (Fig. S2a, Supplemental dataset 8). However, since PSK upregulation was observed around 2 h post wounding, we reasoned that receptor genes might get transcriptionally upregulated after 2 h. RT-qPCR analysis indicated that *OsPSKR4*, *7* and *8* showed a very early induction (Fig. S2b). Although these might contribute in part to wound response, they were unlikely to be involved in wound induced PSK perception which gets triggered in a delayed manner. *OsPSKR5*, *6* and *10* did not show induction post wounding (Fig. S2b). However, we observed a clear and pronounced induction of *OsPSKR*, also known as *OsPSKR12*, which exhibited sharp upregulation 3 h post wounding (Fig. 2c). Interestingly *OsPSKR* was highly upregulated upon OsPep3 treatment and to lesser extent upon OsPep2 treatment and not with other peptides tested (Fig. 2d & Fig. S2a). OsPSKR is a close homolog of AtPSKR1 and DcPSKR, and was previously implicated in insect herbivory responses called as OsLRR-RLK1 (Hu *et al*., 2018). OsPSKR possesses extracellular leucine rich repeats, a single pass transmembrane region and a cytosolic kinase domain (Fig. 2e). Phylogenetic analysis using well-studied ligand binding receptors in *Arabidopsis* and rice indicated that OsPSKR clustered with sulphated peptide-hormone receptors reported from *Arabidopsis*, indicating a possibility that OsPSKR might be involved in perception of PSK which is a sulphated peptide-hormone (Fig. S2c). Homology based model of the protein using AtPSKR1 structure as a reference (Wang *et al*., 2015) was generated to explore if PSK directly interacted with OsPSKR. Docking experiments revealed that PSK potentially interacted with OsPSKR (Fig. 2f). MD simulations at 10 ns were conducted to ascertain protein conformations in the presence of ligand. RMSD plots obtained revealed slightly higher stability of the protein backbone in the presence of ligand (Fig. 2g & 2h, Fig. S2d & S2e). Probable residues taking part in ligand-receptor interactions were deduced from MD simulation experiments with K432 and N433 displaying strong interactions (Fig. 2i). Further, a homology model was built after mutating K432A and N433A. Docking was performed using this mutant and the interactions with PSK were not observed (Fig. 2j). These results suggest that wounding and wound-derived peptide treatment activates PSK signalling that might be perceived by a receptor like kinase, OsPSKR. This signalling is likely to be involved in the activation of downstream responses.

### OsPSKR is a catalytically active kinase that associates with co-receptor OsSERK1

Ligand recognition by LRR-RLKs leads to activation of a pattern recognition receptor (PRR) complex that consists of a cognate receptor, a co-receptor and a receptor like cytoplasmic kinase (RLCK) (DeFalco & Zipfel, 2021). OsPSKR belongs to the subfamily LRR-Xb (Shiu *et al*., 2004) and RD group of RLKs since it has the conserved HRD motif in the catalytic domain. It has been proposed that RD group of kinases function in a catalytic activity-dependent manner (Wang *et al*., 2008; Bender *et al*., 2021). Since OsPSKR possesses a predicted cytosolic kinase domain, we performed an *in vitro* phosphorylation assay to assess its kinase activity. First, we purified the kinase domain (KD) of OsPSKR and the mutant of KD (ΔKD) with mutations in ATP binding site (K814E) and catalytic motif (D912N) (Fig. 3a). We did not notice any autophosphorylation when kinase domain alone was used (Fig. S3a). In many RLKs, it has been shown that the residues undergoing phosphorylation are mostly present in the juxtamembrane (JM) region of the proteins (DeFalco & Zipfel, 2021). Therefore, we considered kinase domain with and without JM region for subsequent assays (Fig. 3a). AtEFR cytosolic domain, which has been shown to undergo autophosphorylation (Bender *et al*., 2021) as well as its kinase dead version were also purified and used as controls in our experiment. We detected phosphorylation signal with the cytosolic domain of OsPSKR and AtEFR, but not in the mutated versions of these proteins suggesting that OsPSKR is a true kinase (Fig. 3b). Most PRRs associate with co-receptors such as AtBAK1 upon ligand binding before activating a series of transphosphorylation events (Ma *et al*., 2016). Rice has two homologs of AtBAK1 namely, OsSERK1 and OsSERK2 with about 79% aa similarity with each other. OsPSKR-GFP or GFP was co-expressed with OsSERK1 or OsSERK2 in *N. benthamiana* to check for receptor-co-receptor associations (Fig. S3b). Co-expression of OsPSKR and OsSERK2 abolished/reduced the accumulation of both the proteins and this was independent of PSK treatment, indicating OsSERK2 was unlikely a co-receptor for OsPSKR (Fig. 3c & Fig. S3c). Therefore, we considered OsSERK1 alone for further experiments. We co-infiltrated OsPSKR-GFP or its kinase domain (KD) mutant (OsPSKRΔKD-GFP) with OsSERK1, followed by super infiltration of PSK. WB analysis revealed that the accumulation of OsPSKR increased in the presence of PSK treatment. This PSK induced stability of OsPSKR was not observed in KD mutant (Fig. 3d & Fig. S3d). PSK induced enhanced accumulation of OsPSKR was also observed in rice plants expressing GFP tagged OsPSKR. This accumulation was more pronounced in the presence of treatment with MG132, a proteasome blocker, indicating a possible proteostatic regulation (Fig. 3e). These results suggest that ligand treatment stabilizes the receptor and this is dependent on the kinase activity of OsPSKR. We further explored if OsPSKR directly interacted with OsSERK1, its likely co-receptor. We co-infiltrated OsPSKR with OsSERK1 or OsSERK2 followed by super infiltration of 1 µM PSK for 20 min. These samples were subjected to immunoprecipitation (IP) of OsPSKR-GFP or GFP. WB analysis indicated the interaction between OsPSKR and OsSERK1, indicating that they are a *bona fide* receptor-co-receptor pair (Fig. 3f). Since IP was performed with the ligand treatment for all the combinations, we further checked the ligand dependency of interaction between OsPSKR and OsSERK1. To test this, we repeated the experiment with or without PSK and observed an interaction between OsSERK1 and OsPSKR independent of ligand treatment. However, endogenous PSK from *N. benthamiana* plants might have played a role in the observed interaction in the absence of exogenous PSK application (Fig. 3g). Nevertheless, these results show that OsPSKR was a catalytically active kinase, and PSK ligand induces its stabilization indicating that PSK was the cognate ligand of OsPSKR and this binding might be required for its strong interaction with its co-receptor.

**Figure 3:**
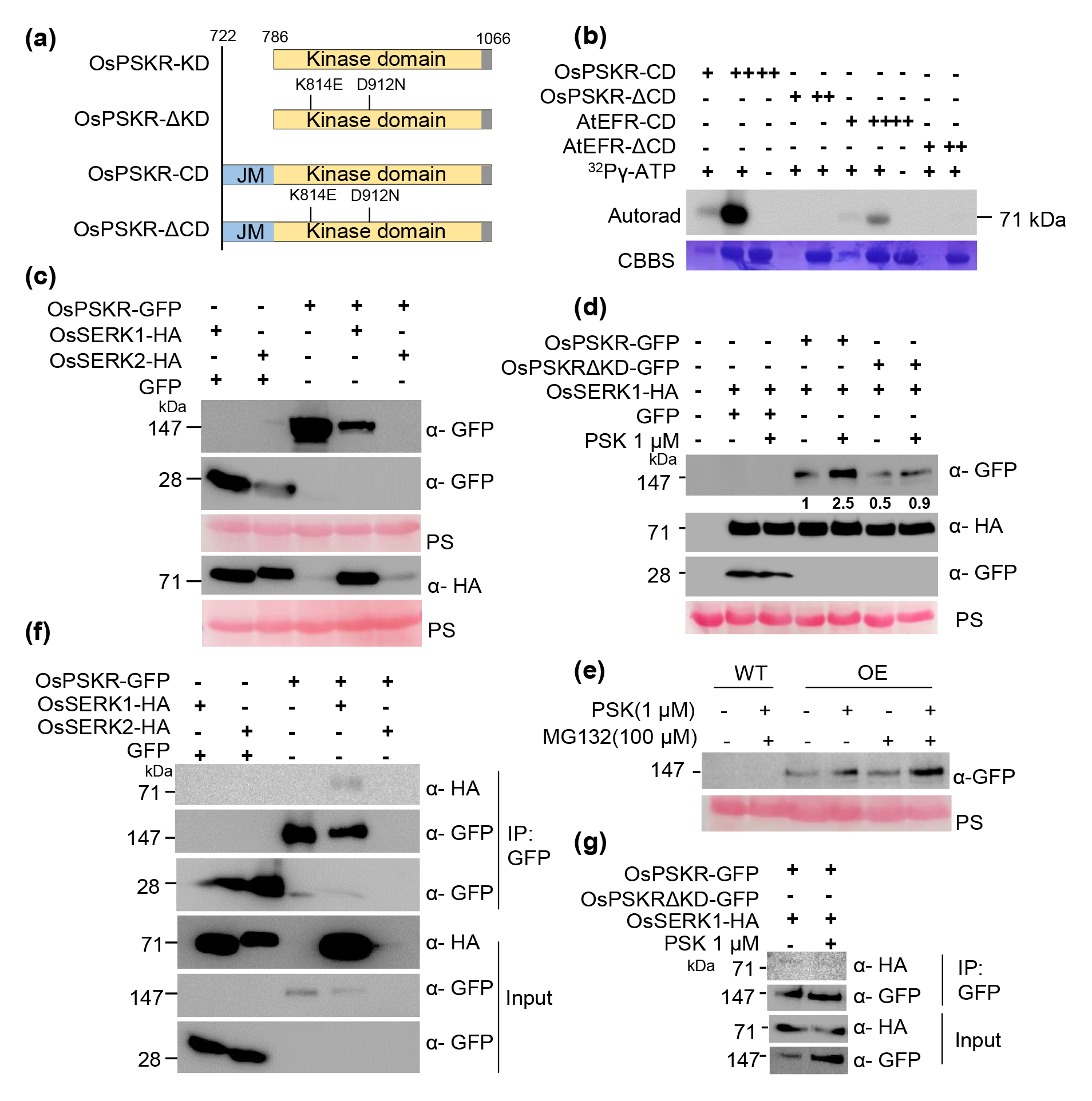
OsPSKR is a catalytically active kinase and associates with OsSERK1. **(a)** Depiction of kinase and JM region of OsPSKR used for kinase assays. KD= Kinase domain, CD-Cytosolic domain. **(b)** OsPSKR is a functional kinase and undergoes autophosphorylation. Autorad-Phosphor signal, CBBS-Coomassie Brilliant Blue. **(c)** Immunoblots showing the accumulation of OsPSKR and OsSERK1/2 in infiltrated *N. benthamiana* plants, PS-Ponceau stain. **(d)** Co-infiltration of OsPSKR/kinase mutant with OsSERK1 in *N. benthamiana* showing ligand dependent increase in WT OsPSKR but not of kinase mutant. **(e)** Western blot showing PSK induced accumulation of OsPSKR in OsPSKR OE rice plants. **(f)** IP assay showing interaction between OsPSKR and SERK1 in the presence of PSK in *N. benthamiana*. **(g)** OsPSKR interaction with OsSERK1 in a ligand independent manner.

### OsPSKR knock-out (KO) plants displayed major growth and reproductive defects

In order to understand the importance of OsPSKR in plant development and wound response in particular, we generated CRISPR-Cas9 mediated KO plants using construct harbouring guide RNA specific to OsPSKR (Fig. S4a). We obtained three independent lines, the nature of mutations in these lines included a two-nucleotide deletion in KO and single nucleotide insertions in both pskr-2 and pskr-3 transgenic lines (Fig. S4b). All these mutations led to premature stop codons with KO-1 terminating at 141 aa and KO-2 and KO-3 terminating at 142 aa. Since these plants had very drastic phenotypes, where the plants failed to make any viable seeds, we also generated artificial miRNA (amiR) mediated knock-down (*kd*) lines (Fig. S4e). amiR that can target OsPSKR, but not other OsPSKRs, designed through WMD3 tool (Schwab *et al*., 2006; Ossowski *et al*., 2008), incorporating position specific GC signatures that improve amiRNA targeting that we deduced previously (Narjala *et al*., 2020) was used for generating transgenic lines. The three *kd* plants obtained exhibited slow growth and reduced fertility, similar to that of KO plants (Fig. 4a and 4b). The expression of amiR was confirmed using northern blotting (Fig. S4f). All *kd* plants had more than 50% reduction of transcript levels exhibiting partial sterility (Fig. S4g). KO plants had reduced height, increased tiller numbers and were completely sterile (Fig. 4a). Surprisingly there were increased number of tillers in KO plants, suggesting a possible compensatory effect (Fig. 4d). The flowers when developed had unusual stigma and style and imperfectly developed palea and lemma (Fig. 4e & i, Fig. S4d). The pollen grains failed to form in KO plants, indicating that reproductive development was completely affected (Fig. 4f & Fig. S4c). Micro-CT and SEM images of individual florets and walls of anthers and stigma (Fig. 4g-h & j) indicated hollow anthers, shrunken anther walls and reduced stigma branching.

**Figure 4:**
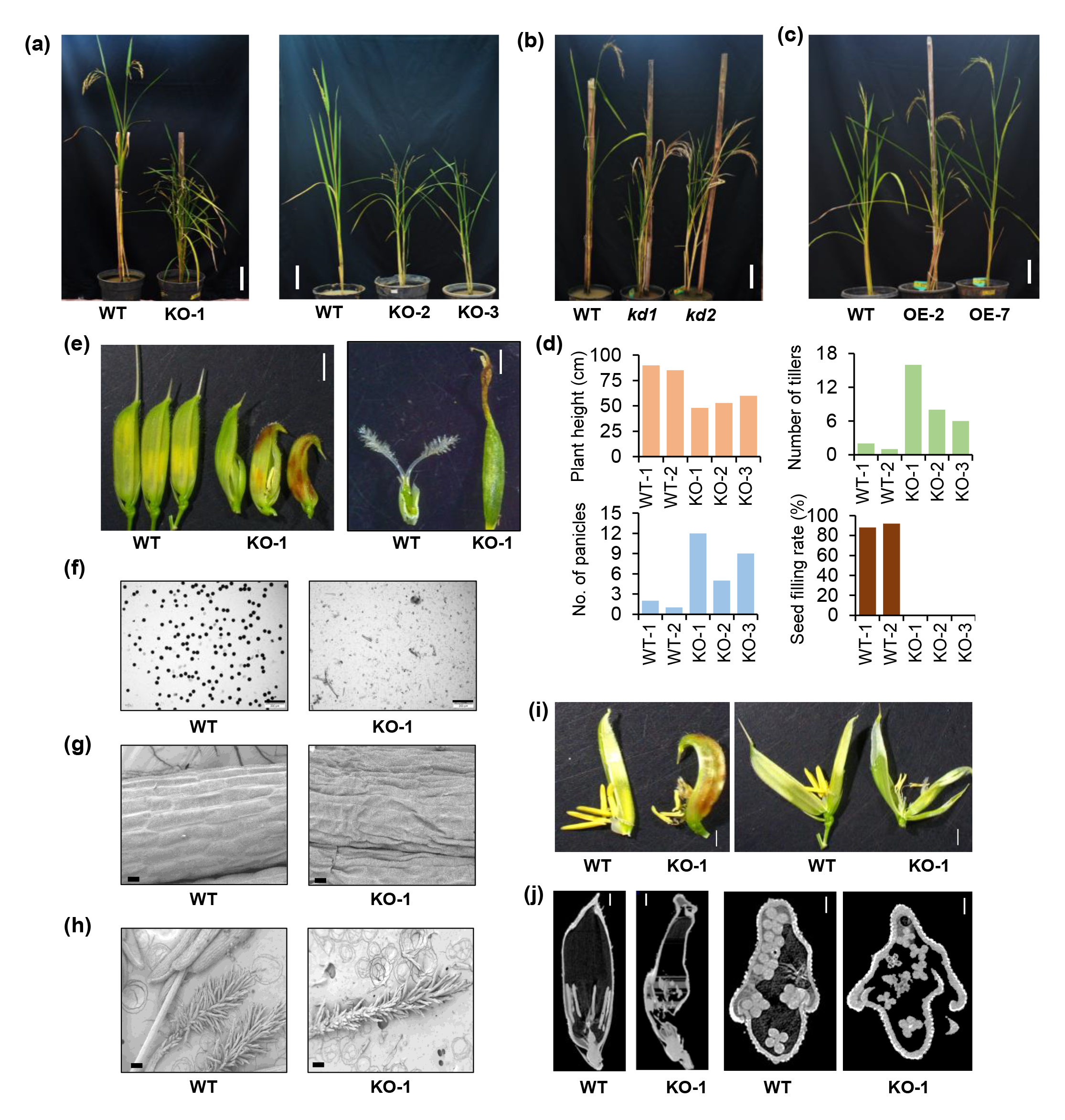
OsPSKR KO plants display major developmental defects. **(a)** Phenotypes of 12-week-old KO plants. scale-8 cm. **(b)** Phenotype of OsPSKR *kd* plants. Scale- 8 cm. **(c)** Phenotype of OsPSKR OE lines. scale-7 cm. **(d)** Measurement of morphological defects in KO plants (Measurements plotted for individual plants). **(e)** KO plants show defective grains and stigma. Scale-0.25 cm for the left-side image and 1 mm for the right-side image. **(f)** Iodine staining of pollen grains in KO plants. Scale-200 µm. SEM images of anther. Scale- 20 µm **(g)** and stigma. Scale- 100 µm **(h)**. **(i)** Anther in KO plants. Scale-0.2 cm. **(j)** Micro-CT images of grains. Scale-800 µm.

We also generated lines overexpressing (OE) OsPSKR under constitutive 35S promoter with GFP tag at the C-terminus (Fig. S5a). We obtained a total of eight plants with two independent T-DNA insertion events with four plants having single copy insertions (Fig. S5b). All plants had increased expression of the transgene (Fig. S5c). Accumulation of OsPSKR-GFP transgenic protein was confirmed using confocal imaging (Fig. S5d), which also indicated membrane localization, expected of pattern recognition receptors. The OE plants clearly showed growth benefits (Fig. 4c). The root growth and shoot growth was better in OE plants compared to wild type indicating the positive regulation of growth by OsPSKR (Fig. S5e & S5f). Together, these results indicated that OsPSKR is an essential gene in rice development, and its involvement in wound response might be crucial for rice plants in countering stresses.

### OsPSKR regulates wound responses

In order to understand the specific roles of OsPSKR in regulation of wound responses and other developmental processes, we performed transcriptome analysis of both KO and OE lines. We identified perturbed expression of several genes in both the cases (Fig. S6a, Supplemental dataset 5). Genes upregulated in KO plants were predominantly involved in purine metabolism and photosynthesis as shown by Gene Ontology (GO) analysis. Downregulated genes in KO included those involved in various metabolic processes including response to wounding category (Fig. S6b). In case of upregulated genes in OsPSKR OE, most were involved in responses to biotic factors and signal transduction. On the other hand, downregulated genes were mainly involved in rRNA processing (Fig. S5c). We further sought to ascertain specific wound response regulation dictated by OsPSKR. Callose deposition is one of the primary responses in case of wounding as well as PTI activation (Vega-Muñoz *et al*., 2020). We assessed callose deposition in KO plants 1 h post wounding by subjecting wounded samples to aniline blue staining.

We observed absence of callose deposits in KO plants indicating that OsPSKR positively regulates wound responses and contributes to callose deposition (Fig. 5a & Fig. S7a). AtPEN3 protein belonging to ABCG transporter class has been attributed to pathogen induced callose deposition in *Arabidopsis* (Matern *et al*., 2019). Expression of rice homologs of AtPEN3 such as *OsABCG37*, *39* and *45* was upregulated upon wounding in WT but either were not upregulated or showed reduced upregulation in KO plants (Fig. 5b), suggesting direct regulation of expression of these genes by OsPSKR.

**Figure 5:**
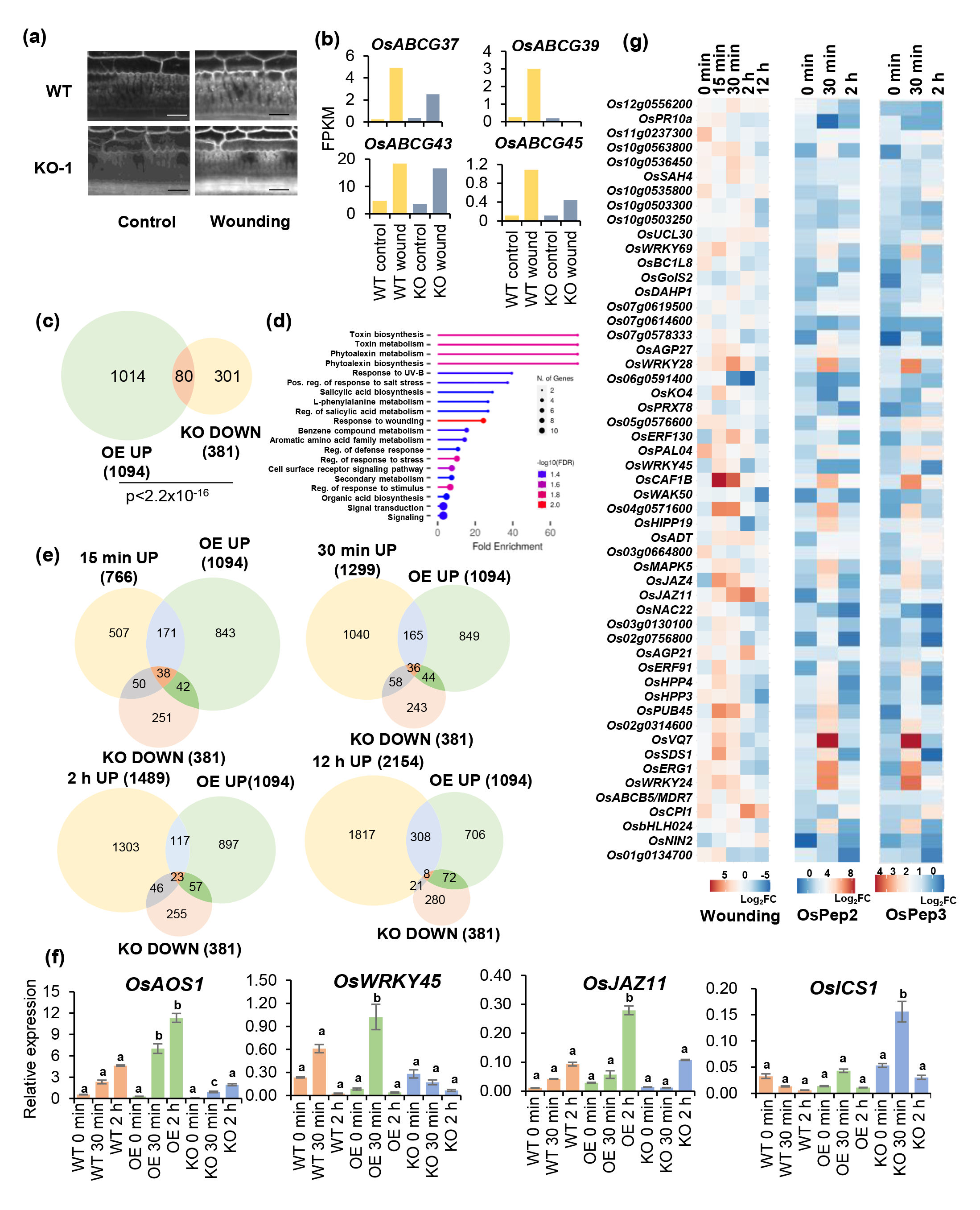
OsPSKR regulates wound responses. **(a)** Callose deposition upon wounding in KO plants. Scale- 25 µm. **(b)** Expression pattern (FPKM values) of ABCG transporters upon wounding in WT and KO plants. **(c)** Venn diagram representing genes overlapping between upregulated genes in OE and downregulated genes in KO. **(d)** GO analysis of genes overlapping between OE and KO lines. **(e)** Representation of overlap between wound responsive genes and genes under OsPSKR regulation across time points. **(f)** RT-qPCR analysis of *OsJAZ11*, *OsWRKY28*, *OsWRKY45* and *OsICS1* relative to *OsActin* upon wounding in WT and transgenic plants, letter codes denote statistical significance (p<0.05), p-values from two-way ANOVA followed by Tukey’s multiple comparison, statistical significance is denoted for OE and KO with respect to WT. Corresponding *p* values are provided in the raw data files. **(g)** Transcriptional dynamics of 53 wound responsive genes under OsPSKR regulation upon wounding and PEP treatment. Fold change for each time-point was calculated with respect to corresponding unwounded/mock treated samples.

To further understand the transcriptional dynamics downstream to OsPSKR which might be part of PSK signalling, we compared the genes that were upregulated in case of OsPSKR OE and the genes that were downregulated in KO plants. We found 80 genes that were common for both these sets, indicating that these genes were under direct regulation of signalling mediated by OsPSKR (Fig. 5c). GO analysis of these 80 genes indicated their involvement in various metabolic processes including wounding (Fig. 5d). We further overlapped the upregulated genes post wounding across time points with genes upregulated in OsPSKR OE and downregulated in KO plants (Fig. 5e). We obtained 53 genes that were unique and were common to all three sets, i.e., upregulated upon wounding across time scales, upregulated in OsPSKR OE lines and downregulated in KO plants. These 53 genes included some of the homologs of well-studied wound responsive genes. These include four WRKY transcription factors- *OsWRKY24*, *OsWRKY28*, *OsWRKY45* & *OsWRKY69*, *OsMAPK5*; ERF transcription factors *OsERF91* & *OsERF130*, two JAZ repressors- *OsJAZ4* & *OsJAZ11*, a VQ domain containing gene- *OsVQ7*, *OsNAC22*; a bHLH transcription factor, *OsCAF1B* and many others (Fig. 5g, Supplemental dataset 6). These results further indicate that OsPSKR is a hub that mediates wound responses in rice. In agreement with this, we observed that the expression of some of the key wound responsive genes was positively impacted in OE and negatively impacted in KO plants upon wounding (Fig. 5f). These genes included *OsAOS1*- involved in JA biosynthesis, *OsWRKY45*- a general stress responsive WRKY transcription factor and *OsJAZ11*- gene involved in repression of JA signalling pathway (Pandey *et al*., 2021). However, *OsICS1* which is not wound responsive in WT plants showed a contrasting trend in KO plants suggesting an antagonistic regulation of *OsICS1* by OsPSKR (Fig. 1c & Fig. 5f). Interestingly, many of the wound responsive genes which were under positive regulation of OsPSKR were also responsive to both OsPep3 and OsPep2 treatments, indicating similarities between signals activated by wounding and wound-derived PEP treatment (Fig. 5g, Supplemental dataset 6). Together, these analyses and results suggest that OsPSKR acts as a positive regulator of early wound responses as observed in callose deposition as well as transcriptional response of some of the early wound responsive genes. Also, OsPeps activated a set of genes similar to those that were under direct regulation of OsPSKR, indicating a conserved and integrated regulation of responses to wounding.

### OsPSKR acts at the intersection of early and late wound responses

KO plants displayed a constitutive cell death phenotype. Cell death and necrosis were more evident near the leaf tips. We performed trypan blue staining and DAB staining of the leaves obtained from all the three KO plants. All three plants displayed constitutive activation of cell death (Fig. 6a). DAB staining revealed exaggerated accumulation of ROS in these plants (Fig. 6b). These phenotypes are suggestive of negative regulation of constitutive defense responses by OsPSKR during homeostatic conditions. In order to understand the regulation in detail, we subjected the cell death displaying leaves from KO for transcriptome profiling. We obtained 1,112 upregulated and 676 downregulated genes including many cell death markers (Supplemental dataset 5). When we intersected the genes upregulated in KO with the genes that were upregulated at 12 h post wounding, we found that over 50% of the genes upregulated in KO plants were also upregulated upon wounding.

**Figure 6:**
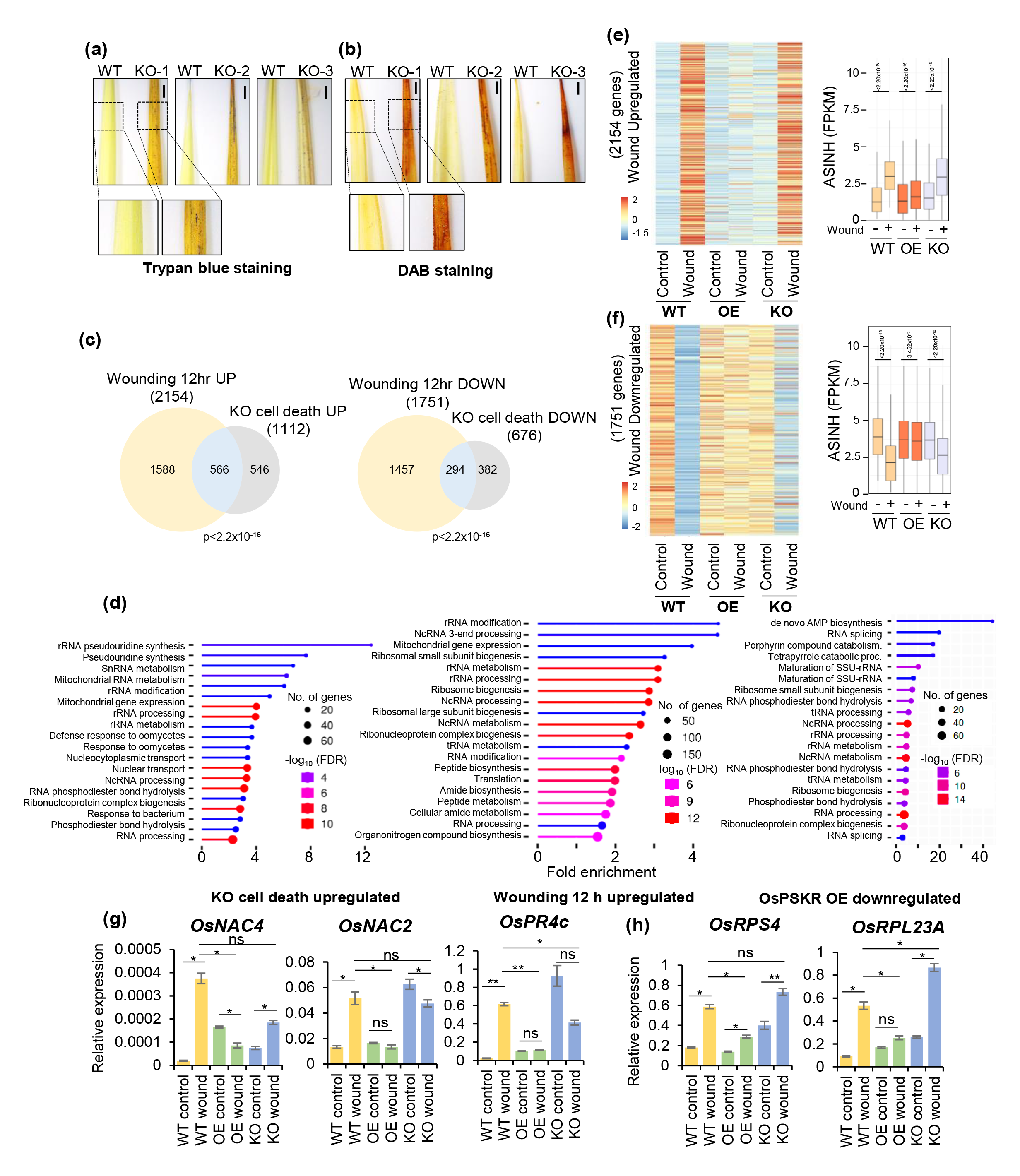
OsPSKR acts at the intersection of early and late wound responses. **(a)** Trypan blue staining indicates constitutive cell death phenotypes in KO plants, scale-1 mm. **(b)** DAB staining for ROS accumulation in KO plants. Scale-1mm. **(c)** Overlap between wound responsive genes and DEGs in KO (Cell-death showing leaves). **(d)** GO analysis of genes upregulated in wounding and KO plants and downregulated in OE. **(e** & **f)** Transcriptional dynamics of wound responsive genes in transgenics, ASINH transformed FPKM values were plotted correspondingly, *p* values obtained from Wilcoxon test are mentioned. **(g)** RT-qPCR analysis of *OsNAC4*, *OsNAC2* and *OsPR4c* post 12 h of wounding relative to *OsActin* in WT and transgenic plants. **(h)** qPCR analysis of *OsRPS4* and *OsRPL23A* post 12 h of wounding in WT and transgenic plants, n=3, pairwise Student’s-*t* test. *: p<0.05, **: p<0.005.

Also, around 50% of the genes which were downregulated in KO were also downregulated upon wounding post 12 h (Fig. 6c). We then performed GO analysis of the genes that were upregulated upon wounding and upregulated in case of cell death displaying KO plants. We observed a great similarity and most of them were involved in rRNA modification and processing (Fig. 6d). Very surprisingly, similar set of genes were downregulated in OE plants (Fig. 6d). These results suggest that OsPSKR maintains the balance between growth and defense signalling during homeostasis and perturbation of the expression of *OsPSKR* leads to perturbation of the balance of signalling. This intersection seems to be at the wound response signalling because KO displayed constitutive upregulation of wound responsive genes and this might be primary cause of cell death observed in case of KO plants. In order to capture the wound responses, we considered all the differentially regulated genes in WT post 12 h. There were 2154 genes that were upregulated in WT, surprisingly same genes did not show any response to wounding in case of OE plants. In KO, there was an exaggerated response for around 50% of the genes indicating that OsPSKR mediated signalling acts as a checkpoint in WT plants and these genes are highly supressed in OE plants (Fig. 6e, Supplemental dataset 7). On the contrary, there were 1751 genes that showed downregulation post wounding in WT. These genes were not downregulated in case of OE and some genes showed a contrasting expression pattern in case of KO plants when compared to WT (Fig. 6f, Supplemental dataset 7). These results collectively suggest that, KO plants display a constitutive upregulation of wound responsive genes that leads to a constitutive cell death phenotype. These wound responsive genes are non-responsive to wounding in OE indicating suppression of wound responses by OsPSKR at later time points post wounding.

Some of the NAC transcription factors have been attributed to hypersensitive responses that lead to cell death phenotype. Increased expression of *OsNAC4* was shown to activate cell death phenotype (Kaneda *et al*., 2009). OsNAC2 was shown to be a regulator of salt-induced cell death (Mao *et al*., 2018). Our transcriptome as well as RT-qPCR analysis showed the upregulation of *OsNAC2* and *OsNAC4* upon wounding in WT plants. The expression of these genes in KO plants upon wounding was similar to that in WT wounded plants. *OsNAC2* was constitutively upregulated in KO plants. However, such a wound-induced upregulation of *OsNAC2* and *OsNAC4* was not observed in OE plants, indicating that OsPSKR suppresses cell death inducers upon wounding (Fig. 6g). *OsPR4c*, a pathogenesis-related gene was upregulated in KO plants and it was wound-responsive in WT plants, and its induction was suppressed in OE plants, suggestive of defense gene suppression upon wounding by OsPSKR (Fig. 6g).

In addition, we observed that ribosomal protein coding genes were upregulated upon wounding in WT plants, and a similar set of genes were upregulated in KO plants displaying constitutive cell death phenotypes. Strikingly, a similar set of genes were suppressed in OE plants suggesting the role of OsPSKR in the regulation of translation. We performed RT-qPCR analysis of genes coding for small and large subunit of ribosomes, *OsRPS4* and *OsRPL23A* respectively. Both of these genes showed upregulation upon wounding and the response was exaggerated in KO and suppressed in OE plants (Fig. 6h). It is possible that OsPSKR mediated signalling culminates in the regulation of genes responsible for translation. These evidences suggest that OsPSKR assists in transitioning of wound responses from early defense responses to late recovery responses. Perturbation of this transitioning leads to exaggerated defense responses which might lead to constitutive cell death which is detrimental to growth. Regulated cell death is key in proper development of reproductive structures and any deviation leads to various developmental defects (Daneva *et al*., 2016).

## Discussion

Plants have evolved a very tightly regulated molecular network in order to respond to wounding and other external cues. Wounding was mostly studied as a part of herbivore responses, or investigated in the context of regeneration responses in which molecular mechanisms leading to effective responses have been discovered (Zhang *et al*., 2019; Radhakrishnan *et al*., 2020; Hernández-Coronado *et al*., 2022; Liang *et al*., 2023). It has been proposed that such mechanisms derived from studies using dicots cannot be generalized for all plants such as monocots having distinct wound induced regeneration responses (Hu *et al*., 2017). Monocot leaves have specialized cells and their growth continues even after mechanical perturbation.

Monocots such as grasses usually fail to acquire stem cell ability and lack regeneration abilities post wounding unlike many dicots (Liang *et al*., 2023). Probing the molecular details involved in wound responses in monocots, in terms of transcriptional changes is important to elucidate distinct mechanisms of wound signalling to ascertain differences from the dicots. The experiments discussed here provide insights into the step-wise molecular signalling events that are a part of the mechanical wounding responses in rice.

Increasing evidences of activation of similar molecular responses upon wounding and insect herbivory, irrespective of insect derivatives, indicated that plants might predominantly rely on endogenously derived signal triggers to activate downstream responses. Therefore, it was important to study wound responses in the absence of foreign (herbivore-derived) cues. Wound perception among dicots involved activation of endogenously derived signalling cues followed by activation of a series of molecular responses leading to active defenses (Vega-Muñoz *et al*., 2020). Our findings indicate the presence of a parallel pathway mediated by an endogenous peptide hormone in signalling wound responses. Very early activation of PEP coding genes upon wounding in rice suggested the importance of PEP induced responses, however, the downstream responses activated by them were not thoroughly investigated. Our multi time-point transcriptome analysis of rice leaves upon OsPep3 treatment revealed the conservation of signal activation between wounding and PEP treatment. The DEGs upon OsPep3 treatment almost completely overlapped with those from OsPep2 treatment indicating the conservation of PEP signalling though the sequences of peptides vary (Fig. S1b). Rice has six PROPEPs and their sequences vary considerably. Bioactive peptides derived from PROPEPs possess a conserved stretch of EGxGGxGGxxH at the C-terminus and usually serve as immune triggers across plants (Fig. S1b). In agreement with this, OsPep3 was able to activate defense signalling suggesting its role as an early defense trigger post wounding. The spatiotemporal regulation of OsPROPEPs, subcellular localization and release into extracellular space upon sensing various cues need to be investigated further to understand if each PEP has a distinct role.

Wound response is a long event and studies show the involvement of various different signalling modules separated temporally (Zhang *et al*., 2019; Hernández-Coronado *et al*., 2022). Many studies in dicots have indicated the presence of a signal relay mediated by phytohormones JA and auxin that promote wound-induced regeneration (Zhang *et al*., 2019; Radhakrishnan *et al*., 2020). Such a signal relay might connect initial defense signals with delayed growth signals. Our transcriptome data indicated the transcriptional upregulation of a precursor coding for another DAMP, namely PSK which is a peptide hormone usually associated with growth signalling (Sauter, 2015). While a majority of the studies implicated JA and auxin hormone mediated wound and regeneration responses, peptide hormone including PSK mediated wound responses have not been thoroughly explored barring in a context-specific transcriptional response (Loivamäki *et al*., 2010).

PSK is a peptide hormone implicated in a multitude of responses (Sauter, 2015) and is perceived by receptor kinases belonging to LRR-RLK family (Matsubayashi *et al*., 2002; Wang *et al*., 2015). In *Arabidopsis* it has been proposed that the cross-talk between brassinosteroid and PSK signalling determines cell fate leading to procambial cell identity (Holzwart *et al*., 2018). Though rice and other monocots lack wound induced regeneration ability, it is possible that the wound induced PSK signalling might contribute to wound induced growth promotion in rice. In rice, 15 PSK receptors were predicted and a few of them were implicated in various stress responses (Nagar *et al*., 2020). Our findings reveal OsPSKR as a novel PSK receptor that was induced post induction of PSK precursor upon wounding. These findings might as well apply to herbivory and other stress responses where PSK might be involved in signalling. OsPSKR was previously implicated in herbivory and was referred to as OsLRR-RLK1 (Hu *et al*., 2018). Other members of the LRR-RLK family that also responded to wounding might perceive other wound-derived signals to initiate parallel responses or they might mediate other stress responses in rice.

**Figure 7:**
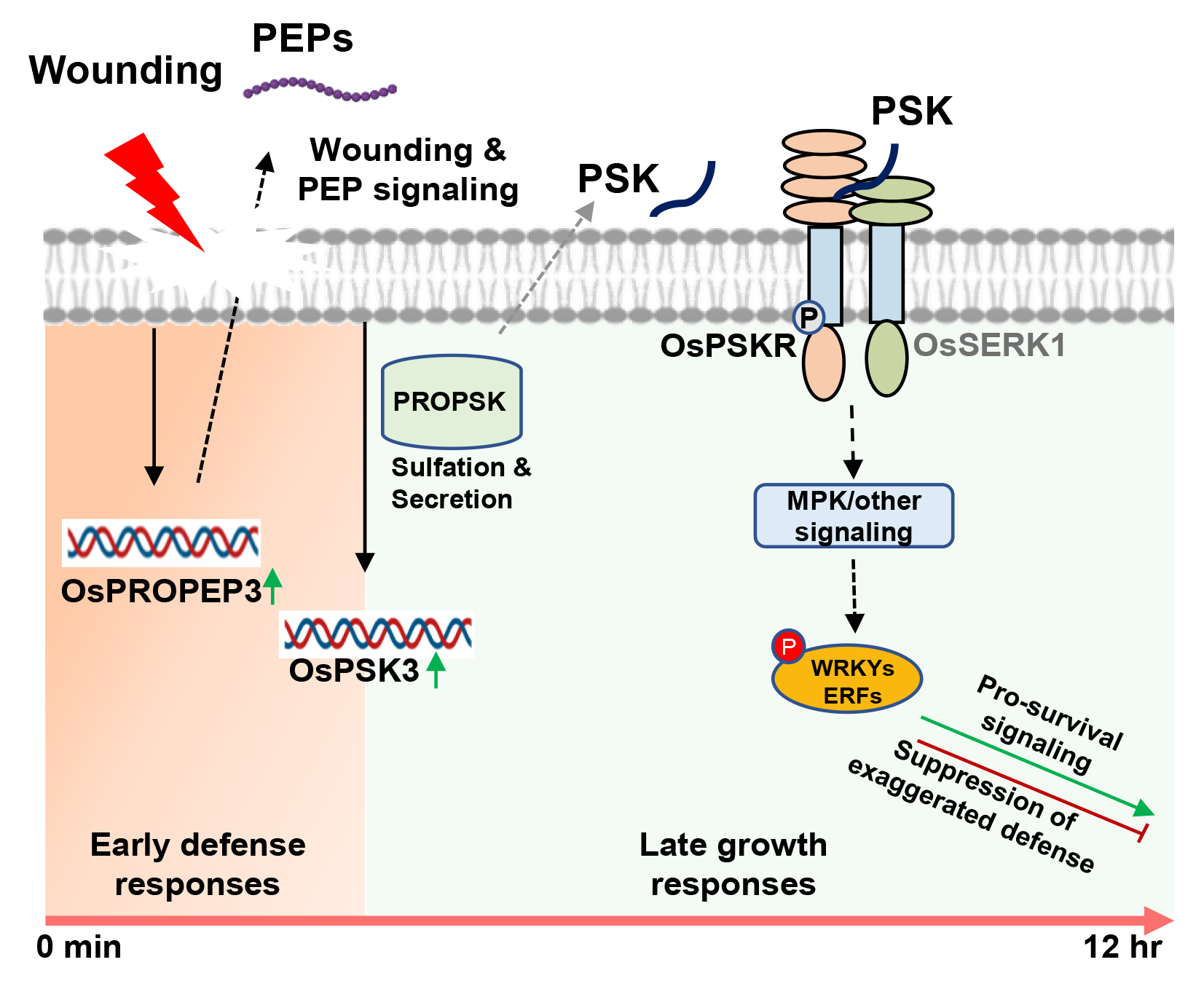
Proposed model for regulation of wound responses by OsPSKR. Wounding triggers the upregulation of precursor of *OsPROPEP3*. Both wound and PEP perception upregulates the expression of precursor of PSK, *OsPSK3*. Upon wounding, PSK signalling gets activated and transduces signal through OsPSKR, receptor of wound-induced PSK. Further, OsPSKR acts at the intersection of transition between defense and growth signalling and acts as pro-growth signal by assisting in suppression of exaggerated defense responses. OsPSKR balances growth and defense responses during homeostasis and activates/represses signals downstream to wound perception.

OsPSKR is an RD kinase and belongs to LRR-Xb subfamily of LRR-RLKs (Shiu *et al*., 2004), members of which are usually part of PRR complexes and activate signalling through a series of transphosphorylation events. We show that OsPSKR is a catalytically active kinase. In agreement with the argument that RD kinases are strong kinases when compared to non-RD kinases (Bender *et al*., 2021), kinase domain of OsPSKR exhibited strong kinase activity in comparison to that of AtEFR KD which is a non-RD kinase (Fig. 3b). OsPSKR also associated with a co-receptor OsSERK1 indicating that signalling activation takes place through interaction between receptor and co-receptor. Increased accumulation of OsPSKR in the presence of ligand indicated the stabilization of protein upon ligand perception. Since this stabilization was not observed in case of kinase activity mutant, it can be perceived that phosphorylation is involved in the stabilization of the protein upon ligand binding. Further studies are required to understand the intricacies of the signalling regulated by OsPSKR and OsSERK1.

PSK signalling has been attributed to growth functions such as tracheary element differentiation, lateral root development, cell fate determination, cell expansion, pollen development, etc. (Matsubayashi & Sakagami, 1996; Matsubayashi *et al*., 1999; Stührwohldt *et al*., 2015; Sauter, 2015; Holzwart *et al*., 2018). However, the loss of function of PSK receptors in other species did not lead to drastic phenotypes unlike what was observed here. Unlike other plants, rice plants compromised in PSK signalling were completely sterile suggesting a strong role of OsPSKR in development. Even the *kd* plants having 50% reduction in *OsPSKR* expression displayed growth defects and partial sterility. On the contrary, the plants overexpressing *OsPSKR* showed growth benefits in terms of plant height and root growth. These observations indicate that PSK signalling and OsPSKR have important functions in rice development beyond wound responses. Interestingly, transcriptome analysis of transgenic plants identified many wound responsive genes under the direct control of OsPSKR mediated signalling. Callose deposition, one of the early responses to wounding is also compromised in the case of KO plants upon wounding, implying the importance of OsPSKR in early wound responses. It is important to note that callose deposition is also a growth and development associated process (Wang *et al*., 2022). In *Arabidopsis*, AtABCG36 which is attributed to callose deposition, regulates transport of both growth and defense related metabolites, resulting in the maintenance of growth-defense balance (Aryal *et al*., 2023). The transport of distinct metabolites by AtABCG36 depends on the phosphorylation status of the protein that is controlled by an LRR-RLK named QSK1 (Aryal *et al*., 2023). Compromised expression of the rice homologs of AtABCG36 upon wounding in KO plants suggested possibly a similar mechanism of regulation of ABCG transporters by OsPSKR either directly or indirectly. In case of OsPSKR OE plants, we observed an increased transcript abundance of genes involved in JA pathway. On the other hand, SA biosynthesis genes were upregulated upon wounding in KO plants, similar to what was previously recorded in a PSKR silenced line upon herbivory (Hu *et al*., 2018). These results are in agreement with the proposed antagonism between JA and SA signalling (Nomoto *et al*., 2021). It is also possible that OsPSKR is playing an important role in the activation of JA signalling as well as in the suppression of SA signalling post wounding.

Unregulated wound responses are detrimental to the fitness of plants and can lead to delayed progression to growth signalling post wounding. KO plants showed constitutive cell death and ROS accumulation phenotypes reminiscent of activation of constitutive wound responses. These phenotypes are suggestive of autoimmune responses and indicate that OsPSKR is a negative regulator of defense responses in homeostasis. It appears that PSK-OsPSKR signalling acts at the nexus of defense and growth signalling that are part of wound responses. Various proteins are being attributed to the regulation of exaggerated defense responses upon wounding. In *Arabidopsis*, BOS1/MYB108 regulates cell death upon wounding and its mutation leads to exaggerated wound induced cell death (Cui et al., 2013). Late wound responses also involve activation of growth and regeneration through signals such as auxin pathway (Zhang et al., 2019). When defense signalling proteins like glutamate receptors were constitutively activated, plant regeneration post wounding was compromised (Hernández-Coronado et al., 2022). A similar mechanism might be present in rice where OsPSKR might assist in promotion of growth signalling post wounding by suppressing defense signalling. In *Arabidopsis*, bleomycin induced cell death was shown to induce AtERF115 in the cells adjacent to treated cells, regulating both regeneration and cell division (Heyman *et al*., 2016). Quiescent cell division mediated by AtERF115 was also dependent on PSK signalling (Heyman *et al*., 2013). We have observed the induction of many ERF transcription factors upon wounding which are also under the positive regulation of OsPSKR. It is possible that wound induced cell death promoted the activity of ERF or related proteins in the adjacent cells that might activate cell division or growth in the neighbouring cells.

PSK signalling contributed to growth mostly by promoting cell elongation in *Arabidopsis* (Kutschmar *et al*., 2009). Wound induced growth promotion and cell death suppression might be regulated by activation of PSK signalling.

Constitutive cell death and ROS accumulation even during homeostasis in KO plants strongly suggested that OsPSKR might be suppressing various defense activators directly or indirectly both at membrane and cytosolic level. Deregulation of AtBAK1, a co-receptor for many ligand binding receptors including AtPSKR1 led to autoimmune phenotypes. This was due to the perturbation of calcium signalling via CNGC19/20 proteins, hyperactivation of NLR proteins and released suppression of other LRR-RLKs (Yu *et al*., 2019, 2023; Wu *et al*., 2020). AtPSKR1 was shown to interact with AtCNGC17 and an ATPase (Ladwig *et al*., 2015). Activation of cell death phenotypes in OsPSKR KO might be due to the perturbation of membrane components that are involved in their dynamics. This possibility needs to be investigated further. Studies in other plant species have showed the role of PSK receptor in attenuation of plant immunity (Igarashi *et al*., 2012). Absence of PSK receptor led to autoimmune responses during beneficial microbe interaction by hyperactivating SA signalling (Song *et al*., 2022). In this study, we observed upregulation of SA biosynthesis gene OsICS1 upon wounding in KO which was otherwise non-responsive to wounding in WT and OE lines, suggesting de-repression of SA signalling upon wounding in KO. Constitutive accumulation of ROS in the KO plants suggested that functional OsPSKR was also involved in maintaining ROS homeostasis. Our findings are in agreement with a recent study that identified the role of PSK signalling in maintaining ROS homeostasis (Hao *et al*., 2023). Since KO plants displayed autoimmune phenotypes, we checked the wound responses in both OE and KO plants. We observed exaggerated wound responses in KO and suppressed wound response in OE indicating that OsPSKR acts at the intersection of defense and growth responses during wounding. Many emerging studies support the role of PSK receptor mediated signalling in growth-defense trade-off. In maize, PSK signalling suppressed cell death during wounding (Koenig *et al*., 2022) similar to the observations reported here. Also, during biotic stress, PSK treatment positively regulated defense responses and suppressed cell death (Zhang *et al*., 2018). We also observed positive regulation of early wound responses and suppression of late wound responses which are responsible for triggering constitutive cell death in KO plants.

This suggests the role of PSK signalling at the nexus of early and late wound responses. PSK signalling also optimized growth-defense through distinct phosphorylation of glutamine synthetase GS2 through a calcium-dependent protein kinase (Ding *et al*., 2023; Förderer & Chai, 2023). Sulphated peptide signalling was attributed to growth-defense trade-off by maintaining growth and defense in the vicinity of damaged cells in *Arabidopsis* (Ogawa-Ohnishi *et al*., 2022). All these studies indicated the importance of sulphated peptide signalling in general and PSK-PSKR signalling in particular in maintaining growth-defense trade-off across various plant species. Our identification of the upstream signalling in PSK signalling, and the downstream events in maintaining growth-defense trade-off upon wounding is timely. Rapidly shifting to growth phase upon wounding might be a strategy evolved by monocots that interacted with herbivores for millions of years. In *Arabidopsis*, the transcriptional activation of PSK and PSKR upon wounding was much delayed (Matsubayashi *et al*., 2006; Loivamäki *et al*., 2010), in comparison to rice as observed here. This suggests a possibility of quick switch to growth signalling post wounding in monocots which might lead to early progression to activate growth than among dicots.

Translational balance is paramount in well-being of all the organisms. Recent studies have shed light on the role of translational perturbation in causing exaggerated cell death in plants (Zhou *et al*., 2023). In KO plants displaying constitutive cell death, we observed increase in the expression of genes involved in translation process. This is reminiscent of constitutive wound responses as the similar set of genes were perturbed upon wounding in WT plants. We also observed that translation related genes were suppressed in OE plants. These results indicate the possible role of OsPSKR in translational regulation. Further studies are required to understand the role of OsPSKR in translational regulation.

Through this study we uncover a very important peptide mediated signal relay responsive to wounding in rice. Such studies can pave way for better understanding of general plant responses that operate by relying on endogenous signals. They also might help in dissecting out the responses that are specific to wounding which are not appreciated in insect herbivore responses. Wounding is not just an adverse reaction akin to pathogenic attack, it also triggers growth and flowering in several horticultural crop plants. Understanding these events and identifying key regulators has implication in improving yield of various crops.

## Methods

### Plant materials and growth conditions

*Oryza sativa indica* PB1 variety was used in this study. Rice plants were grown in greenhouse at 28°C under natural day-night cycle. *N. benthamiana* plants used for infiltration experiments were grown in greenhouse conditions with natural day-night cycle.

### Plasmid construction and cloning

To generate CRISPR knock-out construct, guide RNA fragment was inserted into pRGEB32 vector (Xie *et al*., 2015) using BsaI sites. To generate amiRNA construct, amiRNA sequence was designed using WMD3 (Schwab *et al*., 2006; Ossowski *et al*., 2008) and inserted into pCAMBIA1380 (Genbank accession no. AF234301.1) vector under 35S promoter. To generate 35S:OsPSKR-eGFP (OE plants) plants, CDS of OsPSKR was inserted between SalI and NcoI sites, eGFP coding sequence was inserted between NcoI and SpeI sites of pCAMBIA1380 vector.

For all infiltration experiments derivatives of pCAMBIA1380 were used, 35S promoter sequence was amplified from pRT100 and inserted between EcoRI & BamHI sites. 35S:eGFP was generated by inserting eGFP sequence between BamHI & SalI sites. 35S:OsPSKR was generated as described above. 35S:OsSERK1-HA and 35S:OsSERK2-HA were generated by inserting OsSERK1- HA and OsSERK2-HA sequences between KpnI & SpeI sites in pCAMBIA1380. All vectors were mobilized into GV3101 strain of *Agrobacterium* and confirmed using colony PCR.

### Rice transformation

Rice transformation was carried out as described previously (Sridevi *et al*., 2003; Swetha *et al*., 2018). Rice seeds were dehusked and surface sterilized with 70% ethanol, 4% bleach and 0.1% mercuric chloride. Seeds were placed on callus induction media and kept in dark for 21 days. Calli were used for transformation with the desired constructs using LBA4404 (with pSB1) strain of *A. tumefaciens*. Calli were selected on hygromycin containing media in dark before transferring to regeneration media. Shoots derived from regeneration were transferred to ½ MS media for rooting before transferring to soil.

### Transient expression through *Agrobacterium* infiltration

*A. tumefaciens* strain GV3101 harbouring various constructs were grown in liquid YEB medium with appropriate antibiotics overnight in 28°C shaking incubator as described (Nair *et al*., 2023). Cells were pelleted and resuspended in infiltration buffer containing 10 mM MES (pH 5.7), 10 mM MgCl2 and 100 µM acetosyringone. OD_600_ of the suspension was adjusted to 0.2. Three to four-week-old *N. benthamiana* plants were infiltrated using 1 mL needle-less syringes. Leaves were harvested 3 dpi for immunoblot and IP experiments. For super infiltration of PSK, 1 µM of peptide dissolved in water was used to infiltrate the leaves infiltrated with different *Agrobacterium* cultures containing different constructs after 3 dpi. Leaves were collected 20 min after infiltration unless otherwise mentioned.

### Leaf wounding experiment

Six-weeks-old rice plants were used for wounding. Leaf blades were punched at the edges using a 3 mm punching equipment at a 2 cm interval without damaging midrib (Venu *et al*., 2010). Leaves were collected at corresponding time points along with corresponding unwounded leaves and snap frozen before processing for experiments.

### Peptide treatment

Six-week-old rice leaves were cut into leaf strips of 2 cm each. Leaf strips were maintained overnight in sterile water in six-well plates to wash away the residual wound signals as described previously (Schwessinger *et al*., 2015). Leaf strips were treated with 1 µM of peptides (custom made from Lifetein) and collected at the time points mentioned in the results section with corresponding mock treated samples.

### RT-qPCR analyses

For qPCR analysis, extracted RNA was subjected to cDNA synthesis using Thermo RevertAid RT kit following manufacturer’s protocol. The cDNA obtained was used for qPCR experiments using SYBR green master mix (Solis Biodyne- 5x HOT Firepol Evagreen qPCR master mix). *OsActin* gene primers were used as internal control.

All the primers used are listed in Supplemental Table 1.

### Transcriptome analysis

For RNA sequencing, six-week-old plants were used unless otherwise mentioned. Wounding was carried out as mentioned above and RNA was extracted from leaf samples using Trizol method of RNA extraction procedure. RNA was poly A enriched and Library preparation was carried out using NEBNext® Ultra™ II Directional RNA Library Prep kit (E7765L) according to manufacturer’s instructions. Sequencing was performed in a paired-end fashion on Illumina Hiseq2500 platform. RNA sequence analysis was carried out as described previously (Hari Sundar G *et al*., 2023). An average of 30 million paired-end reads were obtained from an Illumina Hi-seq platform. Adapter trimming was done with Trimmomatic (Bolger *et al*., 2014).

Obtained reads were mapped to rice genome (IRGSP1.0) using HISAT2 (Kim *et al*., 2015). Cufflinks was used to obtain gene expression profiles and results are plotted using R (Trapnell *et al*., 2014). A cut-off of Log_2_FC≥ 1.5 was used for selecting upregulated genes and Log_2_FC≤ −1.5 was used for selecting downregulated genes.

### GO analysis

GO analysis was performed using ShinyGO v0.75 platform. RAPDB gene IDs were used with preference for biological processes with FDR cut-off of P-value:0.05.

### Phylogenetic analysis

Extracellular amino acid sequences of the well-studied ligand binding receptors in *Arabidopsis* and rice were downloaded from TAIR (www.arabidopsis.org) and RAP-DB (https://rapdb.dna.affrc.go.jp/) respectively. Multiple sequence alignment of the receptors was performed using MUSCLE, with their extracellular region under default settings. A neighbor-joining (NJ) phylogenetic tree was generated based on the alignment using MEGA v6.06 (Tamura *et al*., 2013) with the following parameters: Poisson correction and bootstrap values (1000 replicates). The tree was illustrated using iTOL v6 (https://itol.embl.de/).

### Homology modelling and docking studies

Based on sequence similarity and phylogenetic relatedness, a 2.51 Å resolution crystal structure of the AtPSKR1 [PDB ID: 4Z63] was used as template to predict the 3D model of OsPSKR using MODELLER 9v22 (Sali & Blundell, 1993). A set of 10 models were generated for the protein and the protein models with the least DOPE and molPDF scores were selected for final validation. Validation of model quality was carried out using Structure Analysis and Verification Server (SAVES), Protein Quality (ProQ) and Protein Structure Analysis (ProSA) servers. SAVES evaluates the quality of the protein model using various methods such as PROCHECK (Laskowski *et al*., 1993), ERRAT (Colovos & Yeates, 1993) and VERIFY3D (Eisenberg *et al*., 1997).

Structure visualization and analysis were performed using BIOVIA Discovery studio and ChimeraX (Pettersen *et al*., 2021). Ligands retrieved from PubChem and chemspider database for the docking analysis in 3D SDF format were translated and stored in Mol2 format using Open Babel 2.2.3 (O’Boyle *et al*., 2011). The energy minimized modelled OsPSKR structure was processed using the AutoDock tool.

Molecular docking (Ghosh *et al*., 2021) was performed to identify the probable interacting residues between the modelled OsPSKR and the PSK ligand using AutoDock 4.2.1 (Morris *et al*., 2009).

### Molecular dynamics simulation study

Molecular dynamics (MD) simulation studies were carried out as described (Shaw *et al*., 2009) in order to determine the backbone configuration of receptor OsPSKR bound to phytosulfokine peptide ligand. To set up the simulation initially, the systems were built for OsPSKR receptor protein with and without the phytosulfokine ligand in the system builder. MD simulation study was carried out in Desmond vs 2020–1. To set up the initial parameters of an orthorhombic box of 10×10×10 Å, Desmond system builder was used. The receptor OsPSKR and ligand-receptor complex were neutralized with NaCl by adding 0.15 M Na _+_ ions. The prepared systems were relaxed using the Desmond default protocol of relaxation. An MDS run of 10 ns was set up at constant temperature and constant pressure (NPT) for the final production run. The NPT ensemble was set up using the Nosé-Hoover chain coupling scheme at a temperature of 300 K for final production and throughout the dynamics with relaxation time 1 ps. A RESPA integrator was used to calculate the bonding interactions for a time step of 2 fs. All other parameters were associated in the settings followed as described (Shaw *et al*., 2009). After the final production run, the simulation trajectories of OsPSKR receptor complexed with PSK were analysed for the final outcome of RMSD, RMSF, and ligand RMSF, derived from the simulation studies.

### Southern Blotting

Southern blotting was performed as described previously (Pachamuthu *et al*., 2022). Total DNA was extracted from rice using CTAB method (Rogers & Bendich, 1989). Around 10 µg of DNA was used for digestion with the mentioned enzymes. After digestion the DNA was resolved on 0.8% gel in 1XTBE. The DNA was transferred to Zeta probe nylon membrane (Biorad) using capillary transfer method. Membrane was UV crosslinked post transfer. The probe (hygromycin resistance coding DNA) was PCR amplified from the corresponding plasmids used for transformation. The resultant probe was internally labelled with [α-P32] dCTP (BRIT India) using Rediprime labelling kit (GE healthcare) and used for hybridization of the membrane. Further membrane was subjected to washes followed by exposure to phosphor imaging screen and scanned using Typhoon scanner (GE healthcare).

### sRNA northern protocol

Northern hybridization for detecting amiRNA was performed as described earlier (Shivaprasad *et al*., 2012). Around 15 µg of RNA was resolved in a 15% acrylamide gel. Further, electroblotting was performed onto Hybond N+ membrane (GE healthcare). Hybridization was performed using T4 PNK end labelled oligonucleotides with [γ-P32] ATP in hybridization buffer (Ultrahyb buffer-Invitrogen).

### Recombinant protein expression and purification

pGEX-6P-1 (GE healthcare) vector containing Glutathione S-Transferase (GST) tag at the N-terminal end was used for expression of cytosolic domains of OsPSKR and AtEFR. Catalytic residue and ATP binding mutants were generated by site directed mutagenesis (Primers used are listed in the supplemental table 1). The residues mutated for OsPSKR were chosen based on the conservation of residues in other well studied kinases (Hartmann *et al*., 2014). Catalytic mutants for AtEFR were generated based on previously published work (Bender et al., 2021). Rosetta Gami DE3 (Novagen) cells were transformed with pGEX-6P-1 plasmids harbouring cytosolic and mutant cytosolic domains of the above-mentioned proteins. Single colony was used to raise a primary culture of 15 mL in LB broth. The next day, 1% of the primary culture was used to raise a secondary culture of 2 litres and culture was allowed to grow at 37°C until the OD_600_ of 0.5. Further, protein expression was induced by treatment of the culture with 0.3 mM of isopropyl β-D-1- thiogalactopyranoside (IPTG) and allowed to grow at 18°C overnight. Cells were pelleted at 5000 rpm for 1 hr at 4°C and stored at −80°C until use. Further, cells were resuspended in lysis buffer containing 50 mM Tris-HCl pH 8, 300 mM NaCl, 5% (v/v) glycerol, 5 mM β-mercaptoethanol (β-ME), 1 mM phenylmethylsulfonyl fluoride (PMSF), 0.01% IGEPAL, 0.5 mg/mL lysozyme and cOmplete protease inhibitor tablets with EDTA (Roche). Cells were lysed by sonication (65% amplitude, five cycles of 10 seconds pulse-on 15 seconds pulse-off for 2 min) and were subjected to centrifugation at 17,000 rpm at 4°C for 1 h. Supernatant fraction was filtered through 0.45 µm filters. Filtrate was passed through econo-columns (Bio-rad) containing protino glutathione agarose 4B beads (Cytiva) pre-equilibrated with buffer-A (50 mM Tris-HCl pH 8, 300 mM NaCl, 5% (v/v) glycerol, 5 mM β-ME). Flow-through was passed through the column again. Beads in the column were washed by buffer-A and then with buffer-B (50 mM Tris-HCl pH 8, 600 mM NaCl, 5% (v/v) glycerol, 5 mM β-ME) and the again with buffer-A. Protein bound to beads were eluted using 15 mM reduced glutathione prepared in buffer-A. Protein yield and purity was assessed by CBB-staining.

### MPK activation assay

For MPK activation assay, wounded or PEP treated leaf samples were ground using liquid nitrogen. Five volumes of 2x Laemmli sample buffer was added to one volume of ground tissue and boiled at 95°C for 10 minutes as described (Bender *et al*., 2021). Samples were centrifuged and resolved on a 10% SDS-PAGE gel. Proteins were transferred to supported nitrocellulose membrane and probed with p44/42 MAPK antibody from Cell Signaling Technology (CST) (9102) at 1:2000 dilution as described (Bender *et al*., 2021). Bands were detected using Chemiluminescence detector (ImageQuant LAS 4000- GE healthcare).

### *in vitro* protein kinase assay

To check the catalytic kinase activity of OsPSKR, *in vitro* kinase assay was performed as described with slight modification (Bender *et al*., 2021). AtEFR cytosolic domain was used as positive control. About 100 ng for low concentration (+) and 1 µg of protein for higher concentration (++) were used. Proteins were incubated in 20 µL kinase buffer (50 mM Tris-HCl pH 7.4, 100 mM NaCl, 2.5 mM MgCl_2_, 2.5 mM MnCl_2_, 10 µM ATP, 5% (v/v) glycerol, 370 kBq γ^32^P-ATP) at 30°C for 30 min. Reaction was stopped by adding 20 µL 2X Laemelli SDS loading buffer and boiling at 70°C for 5 min. Proteins were separated by SDS-PAGE and further exposed to phosphor-screen for 10 min. Exposed screen was scanned and imaged using molecular imager (GE).

### Protein extraction, immunoblotting and IP

The samples were ground using mortar and pestle and around 200 mg tissue was taken for protein extraction as described previously (Yang *et al*., 2022). Ground tissue was resuspended in 200 µL extraction buffer (100mM Tris-HCl pH 7.5, 1 mM EDTA, 150 mM NaCl, 1% Triton X-100, 0.1% SDS, 10 mM dithiothreitol (DTT) and 1x protease inhibitor cocktail). Equal amount of 6x SDS-PAGE loading buffer was added and the samples were boiled at 95°C for 10 min, and then the samples were centrifuged at 13000 rpm for 10 min. Supernatant was collected in a fresh tubes and samples were resolved on 8% SDS-PAGE gels. Proteins were transferred to nitrocellulose membrane. Membrane was blocked with 5% milk and probed with primary antibodies anti-GFP (Sigma) and anti-HA (CST). Further, the membranes were probed with anti-rabbit secondary antibody and imaged using image quant. Same procedure was followed for protein extraction from rice leaves except that 1% (w/v) PVP was added to the extraction buffer.

For IP, leaves of infiltrated *N. benthamiana* plants were harvested at 3 dpi. For rice, six-week-old plants were used. IP was performed as described previously (Nair *et al*., 2020). Around 2.5 gm of ground tissue was resuspended in 3 mL of extraction buffer (50 mM Tris-HCl pH 7.5, 150 mM NaCl, 2 mM EDTA, 10% glycerol, 2 mM DTT, 2 mM Phenylmethanesulfonyl Fluoride (PMSF), 1 % Triton-X, 1x protease inhibitor cocktail) and homogenised for 15 min at 4°C. Further, additional 9 mL of extraction buffer without Triton-X was added and homogenised for 30 min at 4°C. Samples were centrifuged at 5000 rpm at 4°C and supernatant was passed through 0.45 µM cell strainers. Then, 30 µL of GFP-trap (Chromotek) beads were added to each sample and binding was allowed for 3 h at 4°C with slow rotation on roto-spin. After binding, beads were collected using magnetic stand and subjected to three washes with wash buffer (50 mM Tris-HCl pH 7.5, 150 mM NaCl, 1 mM PMSF). Proteins were eluted from the beads by boiling with 50 µL of 2x SDS buffer at 80°C. Proteins were resolved on 8% SDS-PAGE gels and detected using WB.

### SEM imaging and micro-computed tomography (Micro CT)

SEM imaging was carried out as described previously (Pachamuthu *et al*., 2021). Rice spikelets were collected prior to flowering and were fixed in 16% formaldehyde, 25% glutaraldehyde and 0.2M cacodylate buffer for 12-16 h. Further, samples were washed with double distilled water and subjected to series of dehydration using series of 25-100% ethanol. Samples were then dried using Leica EM CPD300, gold coated, and imaged using a Carl Zeiss scanning electron microscope at an accelerating voltage of 2kV.

### Pollen staining

Pollen grains viability test was performed using I_2_-KI staining solution containing 0.2% (w/v) I_2_ and 2% (w/v) KI as described (Pachamuthu *et al*., 2021). Anthers from (n=6) spikelets of mature two individual panicles just one day before the fertilization were collected in 100 µl of I_2_-KI solution. Pollen grains were released in the solution by mechanical shearing using micro tips. After 10 min viable pollen grains in 20 µl solution were counted under bright-field microscope (Olympus BX43). Round and dark blue were considered as viable pollen grains, while light blue or distorted shape pollens were considered as nonviable.

### DAB staining

DAB staining procedure was performed as described previously (Melvin *et al*., 2017). Leaves were immersed in DAB staining solution (dissolved 1 mg/mL of DAB by reducing the pH to 0.3 using 0.2 M HCl, further added 0.05% (v/v) tween20 and 200 mM Na_2_HPO_4_ to obtain 10 mM Na_2_HPO_4_ DAB staining solution) overnight on a rotor. Samples were bleached using bleaching solution (ethanol: acetic acid: glycerol – 3:1:1) and washed thrice for 10 min each with boiling.

### Trypan blue staining

Trypan blue staining solution (for 40 mL – 10 mL of lactic acid (85% w/w), 10 mL of phenol (TE buffer equilibrated, pH 7.5-8.0), 10 mL of glycerol, 10 mL of distilled water, 1 mg/mL of trypan blue) was prepared and leaf samples were stained for 1 hr as described (Yu *et al*., 2019). Samples were destained by giving repeated washes with 100% ethanol.

### Callose staining and imaging

About 0.01% of aniline blue in KPO_4_ buffer pH-7.5 was used for staining leaf samples. Staining was performed at room temperature overnight. Stained samples were imaged using confocal microscopy (Carl Zeiss LSM980). Excitation wavelength of 420-480 nm and emission wavelength of 495-550 was used as described previously (Scalschi *et al*., 2015).

## ACKNOWLEDGEMENTS

We thank members of Shivaprasad lab for suggestions. We thank the Next Generation Genomics, radiation, CIFF and EM at NCBS-TIFR, Bangalore. We thank Prof. K. Veluthambi for binary vectors, PB1 seeds and *Agrobacterium* strains. This study was supported by Department of Atomic Energy, Government of India, under Project Identification No. RTI 4006 (1303/3/2019/R&D-II/DAE/4749 dated 16.7.2020). This work was also supported by grant BT/PR12394/AGIII/103/891/2014;BT/IN/Swiss/47/JGK/2018-19; from Department of Biotechnology, and MST/PRAO/Control/2020-21/PFMS from Department of Science and Technology, Government of India.

## AVAILAIBLITY OF DATA AND MATERIALS

All data generated or analysed during this study are included in this published article (and its supplementary information files). Source data for all the images, gels and blots used in the study are provided as original raw file, and images or blots used in the figure files are also provided as uncropped images with relevant area labelled.

RNA-Seq data is available under GEO accession number: GSExxxxxx

## AUTHOR CONTRIBUTIONS

CYH and PVS conceptualized the study, analysed the data and wrote the manuscript. CYH performed most of the experiments. AP performed SEM imaging. AN helped in kinase assay. MC performed modelling and docking. SR performed confocal imaging.

## Supplemental information

Manuscript has 7 Supplemental Figures, 2 Supplemental Tables and 8 supplemental datasets.

### Supplemental figures

**Figure S1:**
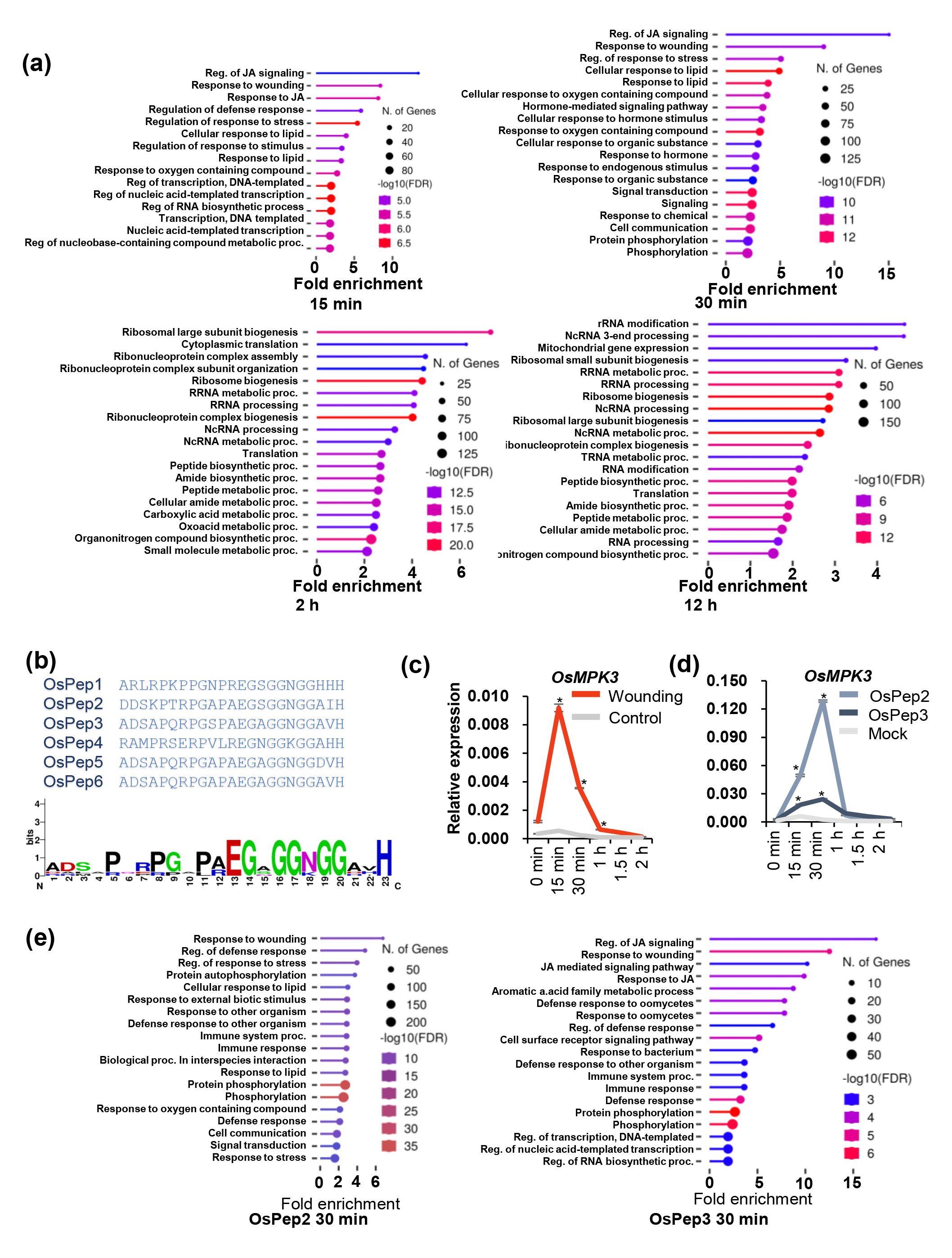
PEP treatment triggers similar set of genes that are involved in wound responses. **(a)** GO analysis of significantly upregulated genes upon wounding across different time points, Log2FC>1.5, p<0.05. **(b)** The sequences of bioactive PEPs derived from their precursor PROPEPs. **(c)** RT**-**qPCR analysis of expression of *OsMPK3* upon wounding. **(d)** qPCR analysis of expression of *OsMPK3* upon OsPep2 and OsPep3 treatments. **(e)** GO analysis of upregulated genes upon peptide treatment at 30 min post treatment, Log2FC>1.5, p<0.05. (c) and (d): n=3, pairwise Student’s-*t* test, *: p<0.05.

**Figure S2:**
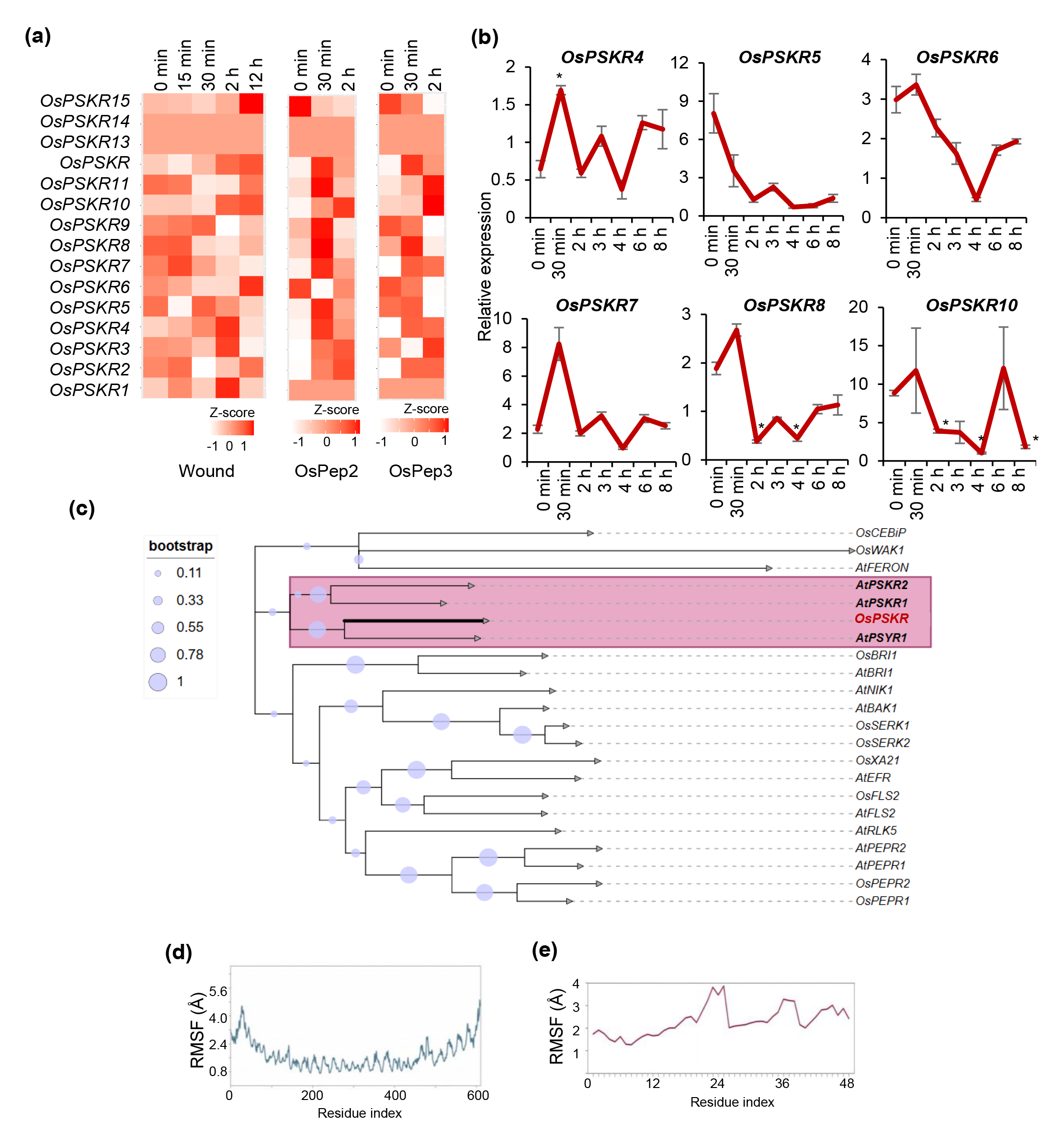
OsPSKR is a wound-responsive PSK receptor. **(a)** Expression profile of all predicted OsPSKRs upon wounding and peptide treatment. **(b)** qPCR showing expression dynamics of possible wound responsive *OsPSKRs*, n=3, pairwise t-test, * = p<0.05. **(c)** Phylogenetic analysis of well-studied plant LRR-RLKs. **(d)** RMSF plot showing conformation of OsPSKR. **(f)** RMSF plot of ligand.

**Figure S3:**
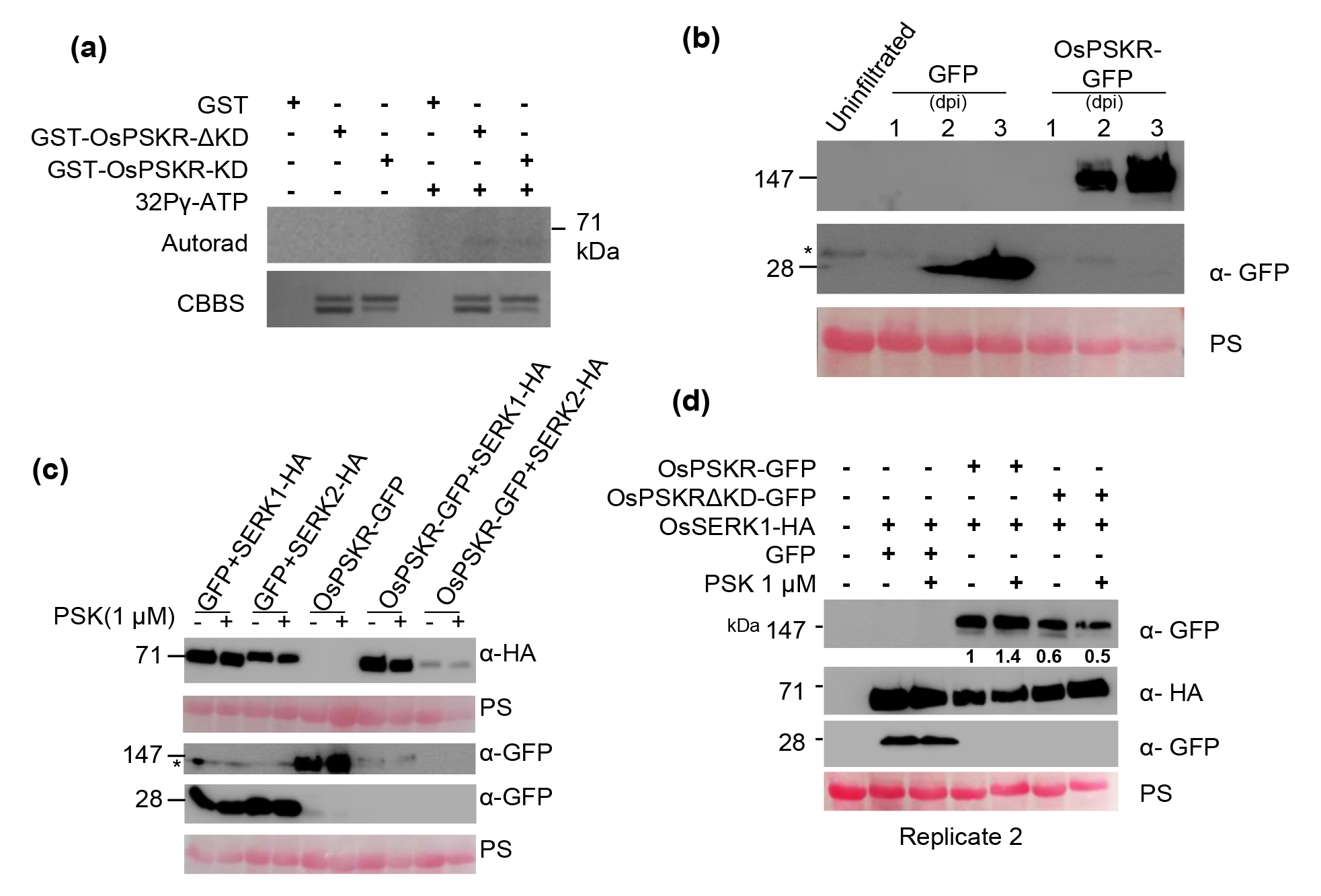
OsPSKR requires JM region for autophosphorylation and ligand induced accumulation of the protein is kinase activity dependent. (a) Autophosphorylation of OsPSKR requires JM region. (b) Transient expression of OsPSKR in N. benthamiana. (c) Co-infiltration of OsPSKR and OsSERK1/2. (d) Co-infiltration of OsPSKR/kinase mutant with OsSERK1. *= non-specific bands.

**Figure S4:**
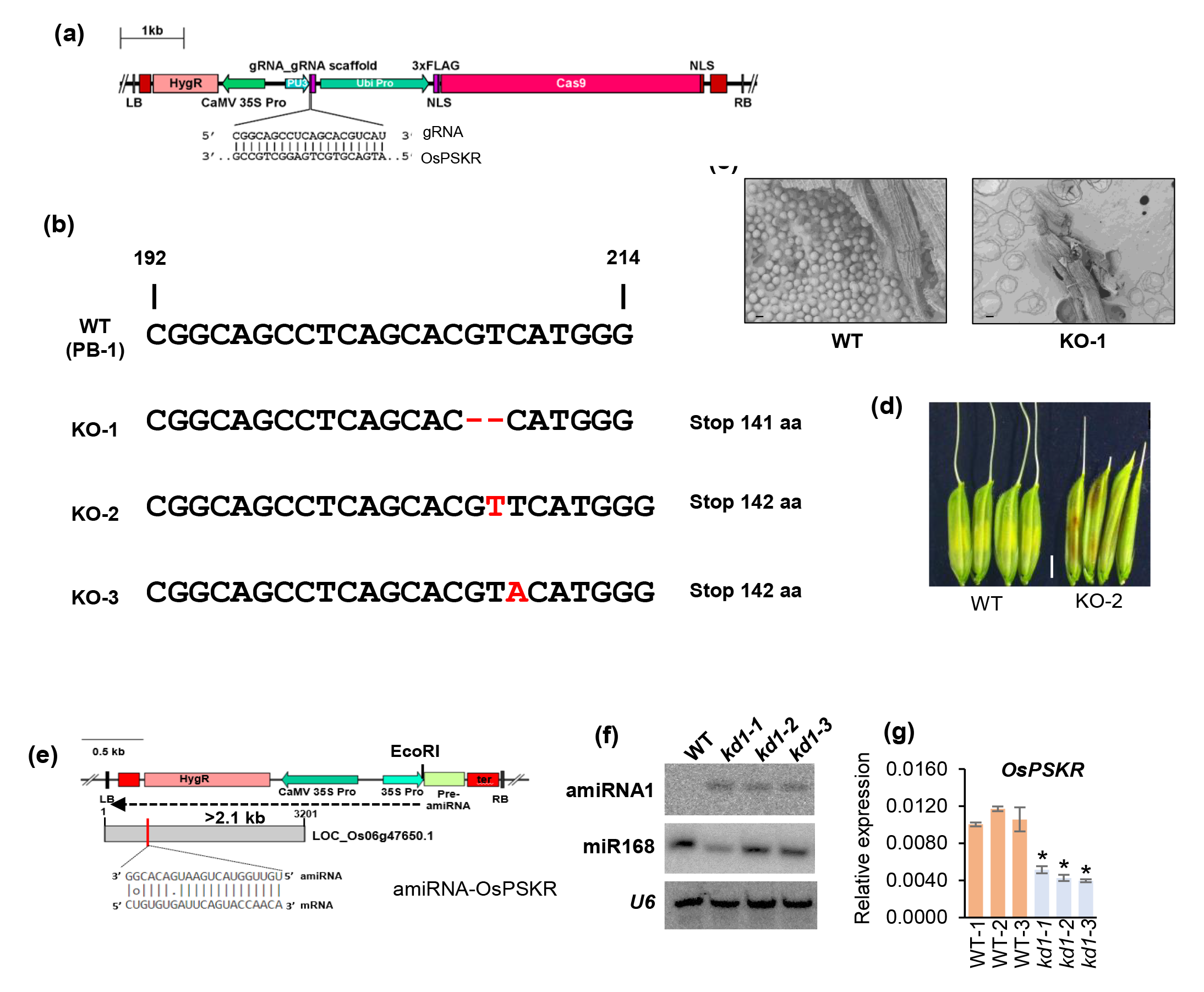
OsPSKR KO and kd plants display similar phenotypes. **(a)** Linear T-DNA map depicting gRNA construct used for CRISPR KO of OsPSKR. **(b)** Nature of mutations in KO plants. **(c)** SEM images of pollen grains in KO plants. scale-20 µm **(d)** Grain morphology is compromised in KO-2 plant as well, scale- 0.25 cm. **(e)** Linear T-DNA map depicting amiRNA construct used for *kd* of OsPSKR. **(f)** Northern blot showing the presence of amiRNA-OsPSKR. (**g)** qPCR analysis of expression of *OsPSKR* relative to *OsActin* in *kd* plants, n =3, student’s t-test compared to WT-1, * represents p value<0.05.

**Figure S5:**
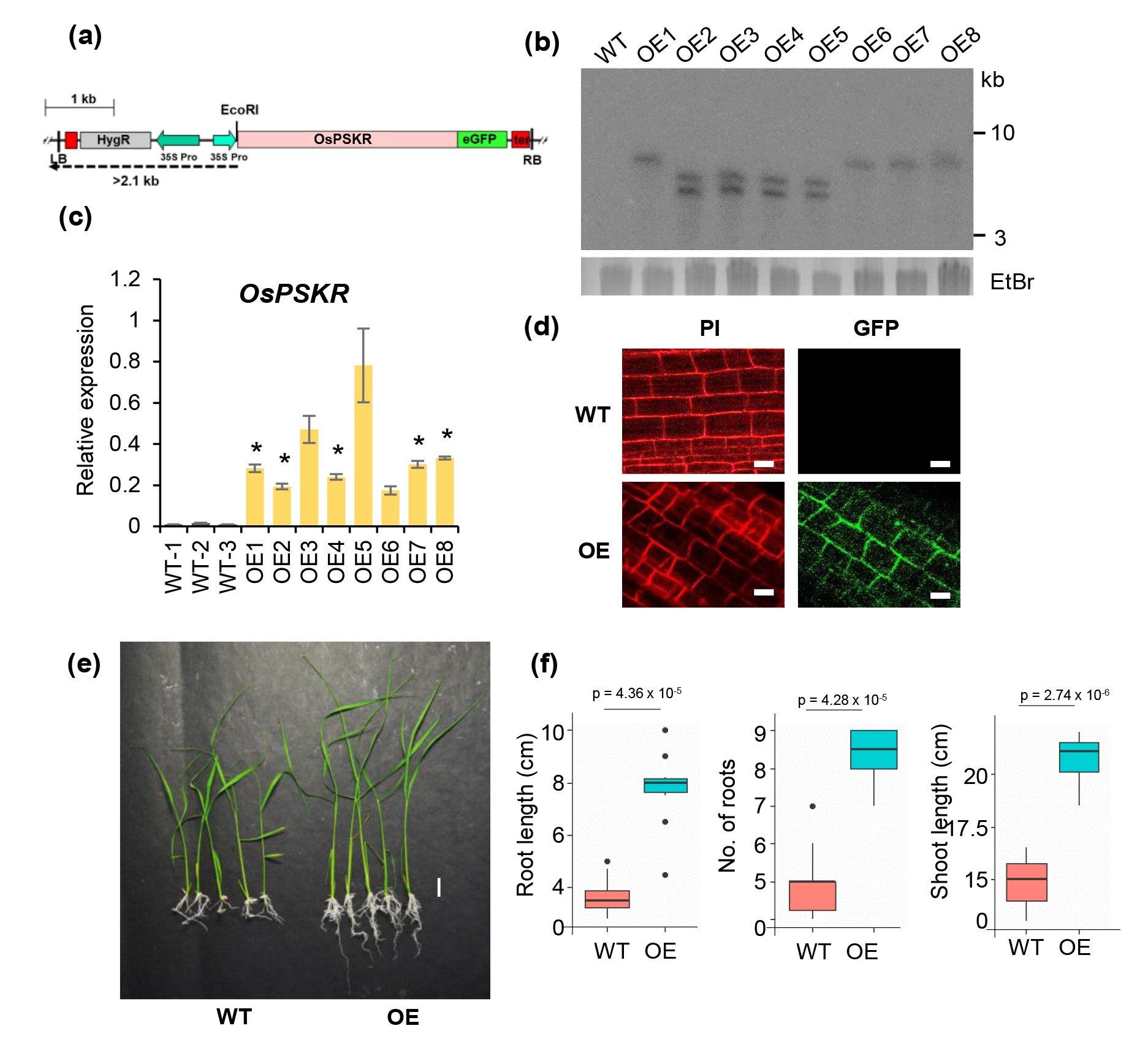
Validation of OsPSKR OE. **(a)** Linear T-DNA map depicting OsPSKR OE under 35S promoter. **(b)** Southern blot showing junction fragments in OE lines. **(c)** qPCR analysis of expression of *OsPSKR* relative to *OsActin* showing OE in transgenic plants, n =3, student’s t-test, * represents p value<0.05. **(d)** Confocal images of roots displaying membrane localization of OsPSKR. Scale-39 µm. **(e)** Phenotype of 3-week-old seedlings of WT and OE plants. Scale-1 cm. (f) Phenotype measurements of OE seedlings compared to WT, n=10, pairwise Student’s-*t* test.

**Figure S6:**
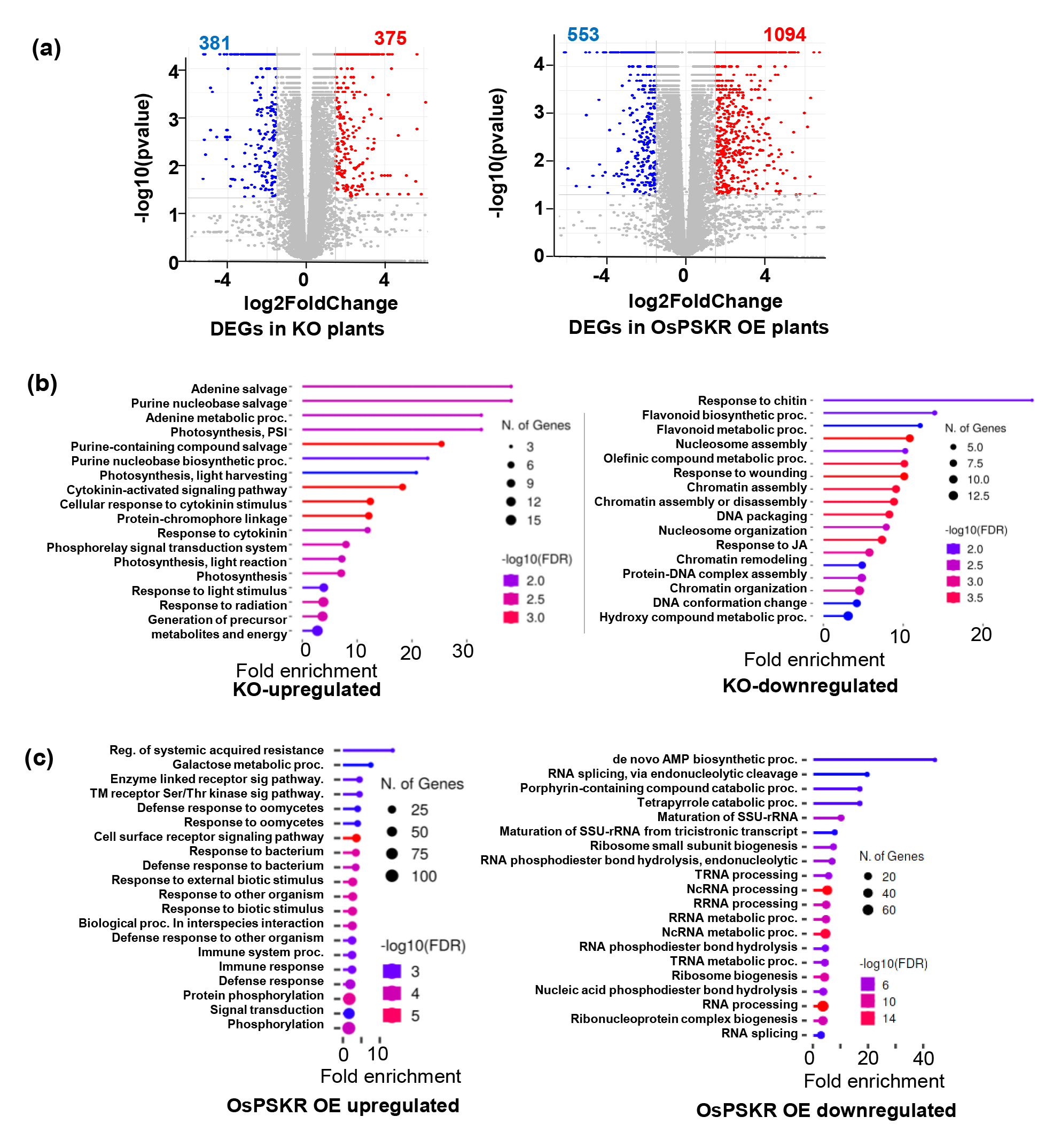
Differential expression analysis of genes in KO and OsPSKR OE plants. **(a)** Volcano plot representing differentially expressed genes in KO and OsPSKR OE plants. Blue represents downregulated (Log2FC ≤ −1.5) and red represents upregulated (Log2FC ≥ 1.5). **(b)** GO analysis of differentially expressed genes in KO plants. **(c)** GO analysis of differentially expressed genes in OsPSKR OE. Log_2_FC ≥1.5 for upregulated & Log_2_FC ≤-1.5 for downregulated, p<0.05.

**Figure S7:**
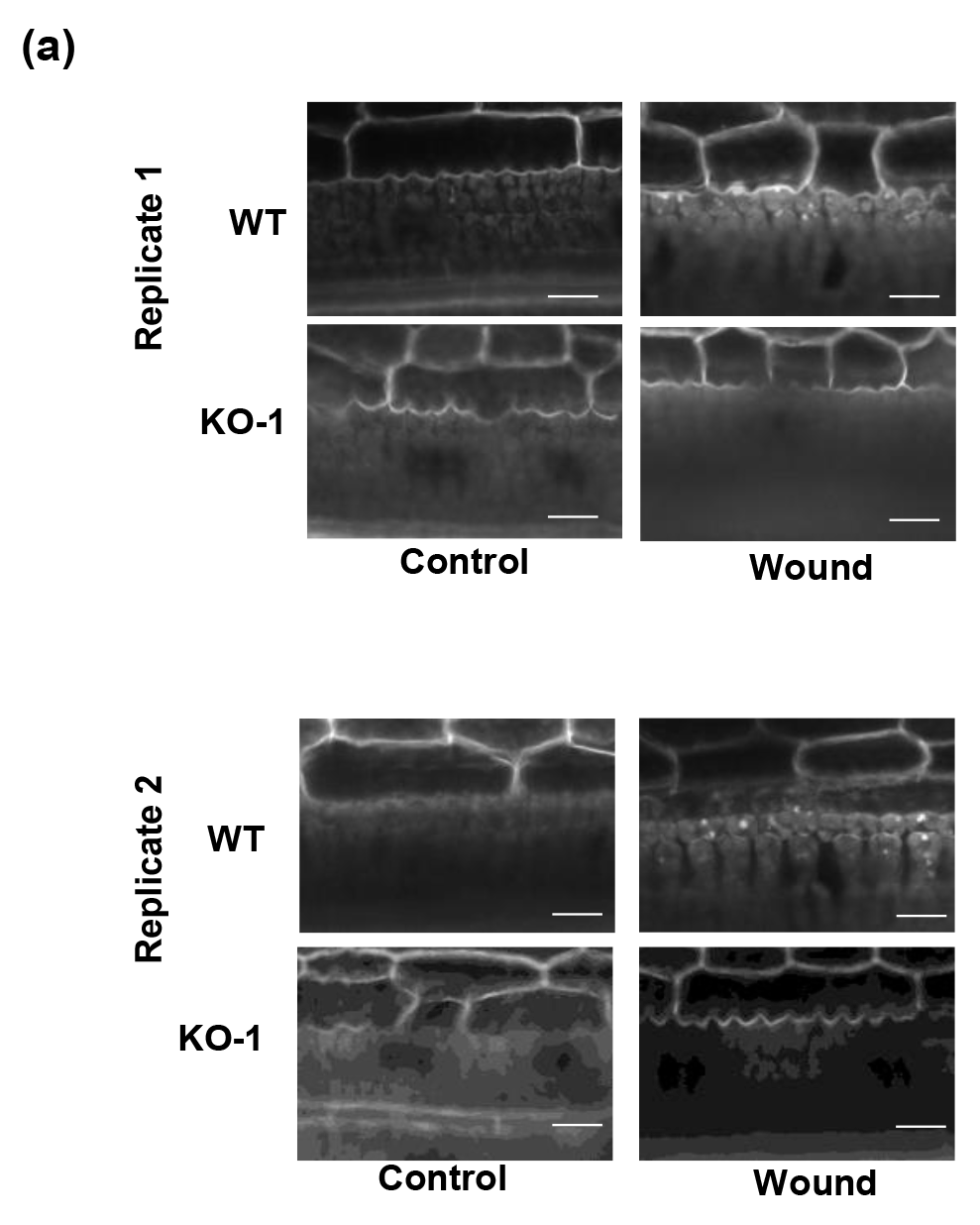
OsPSKR positively regulates callose deposition upon wounding. (a) Callose deposition upon wounding (1 h) in WT and KO leaves. Scale- 25 µm.

### Supplemental Tables

**Table S1:**
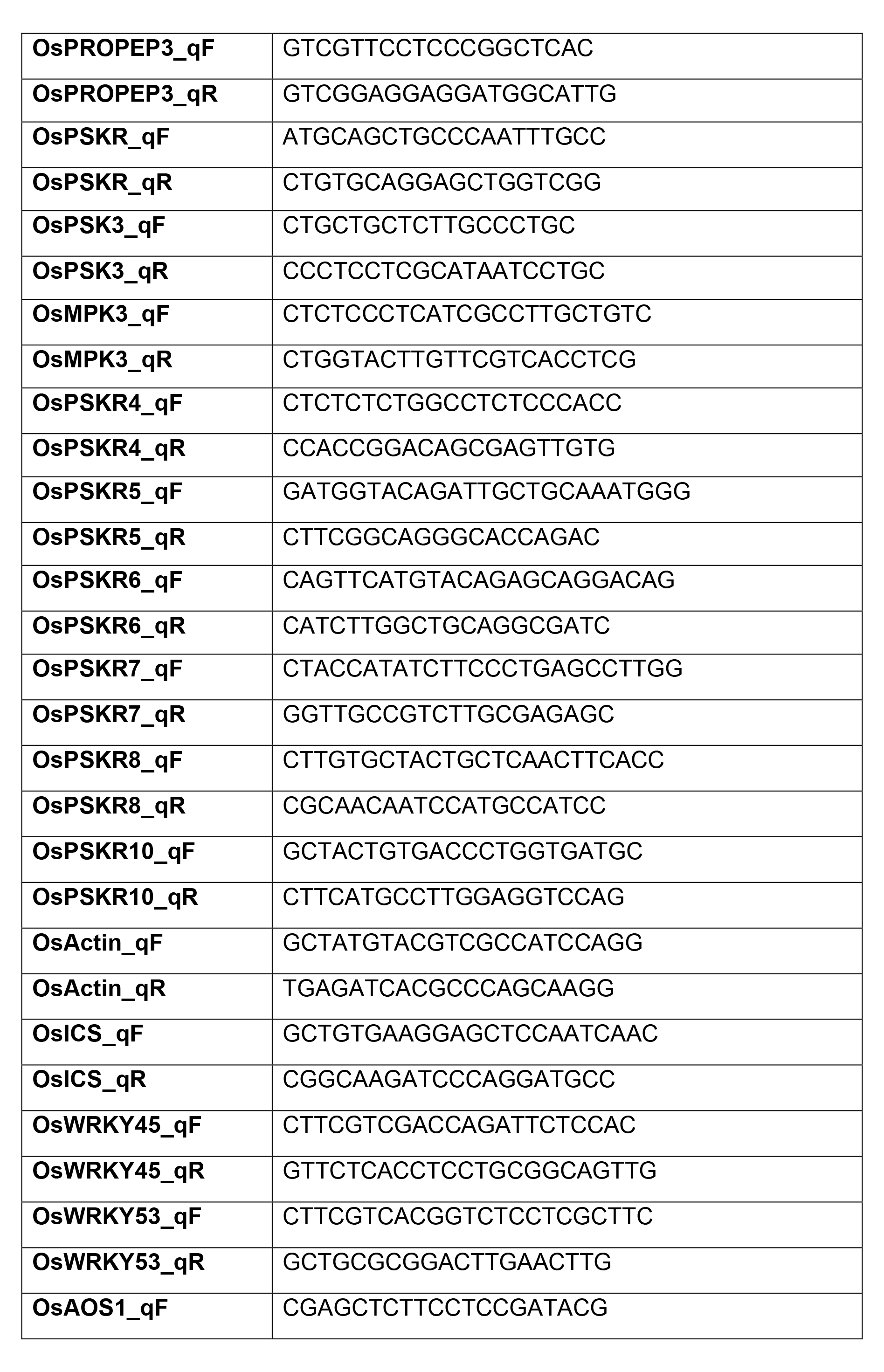
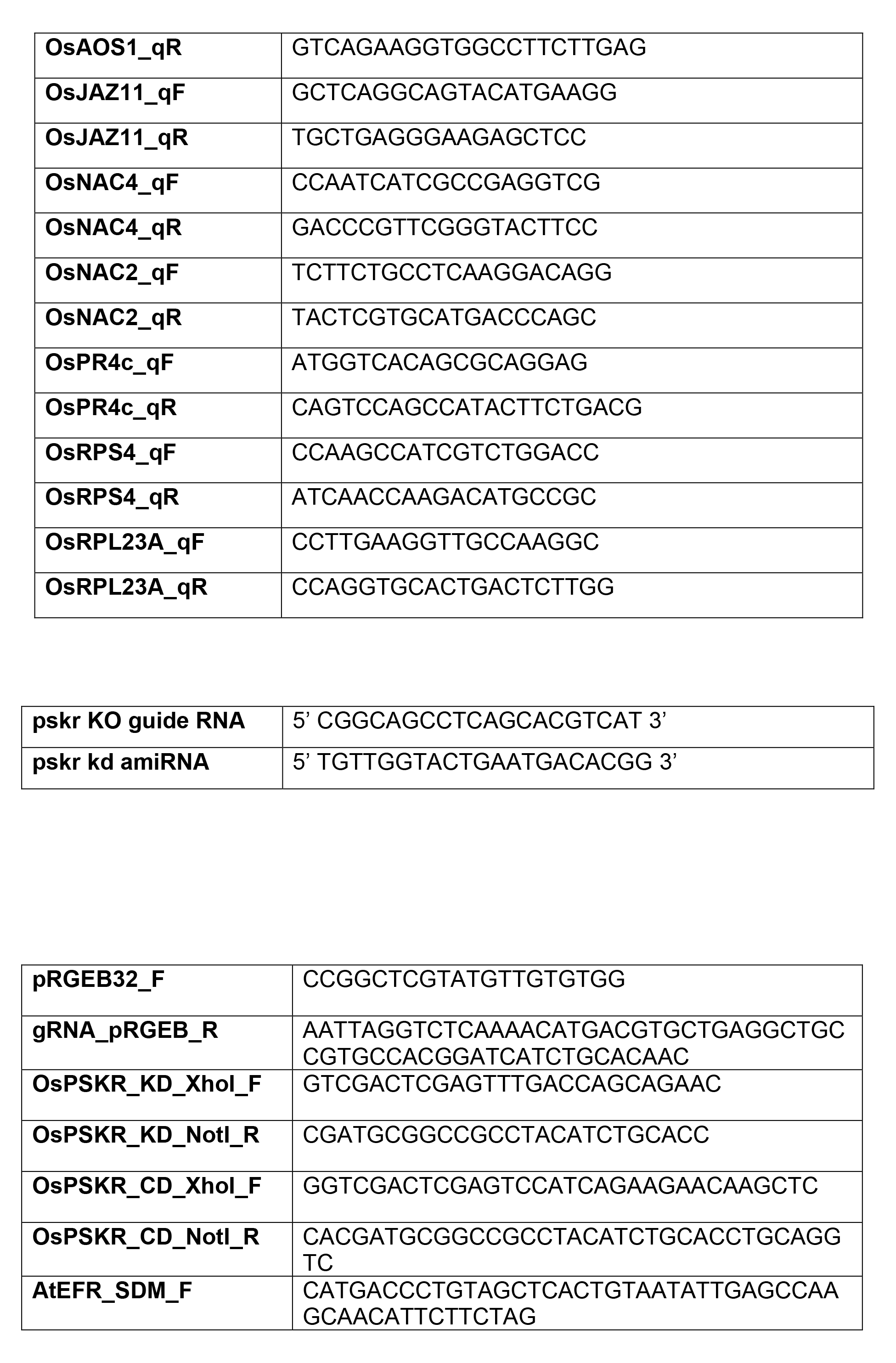
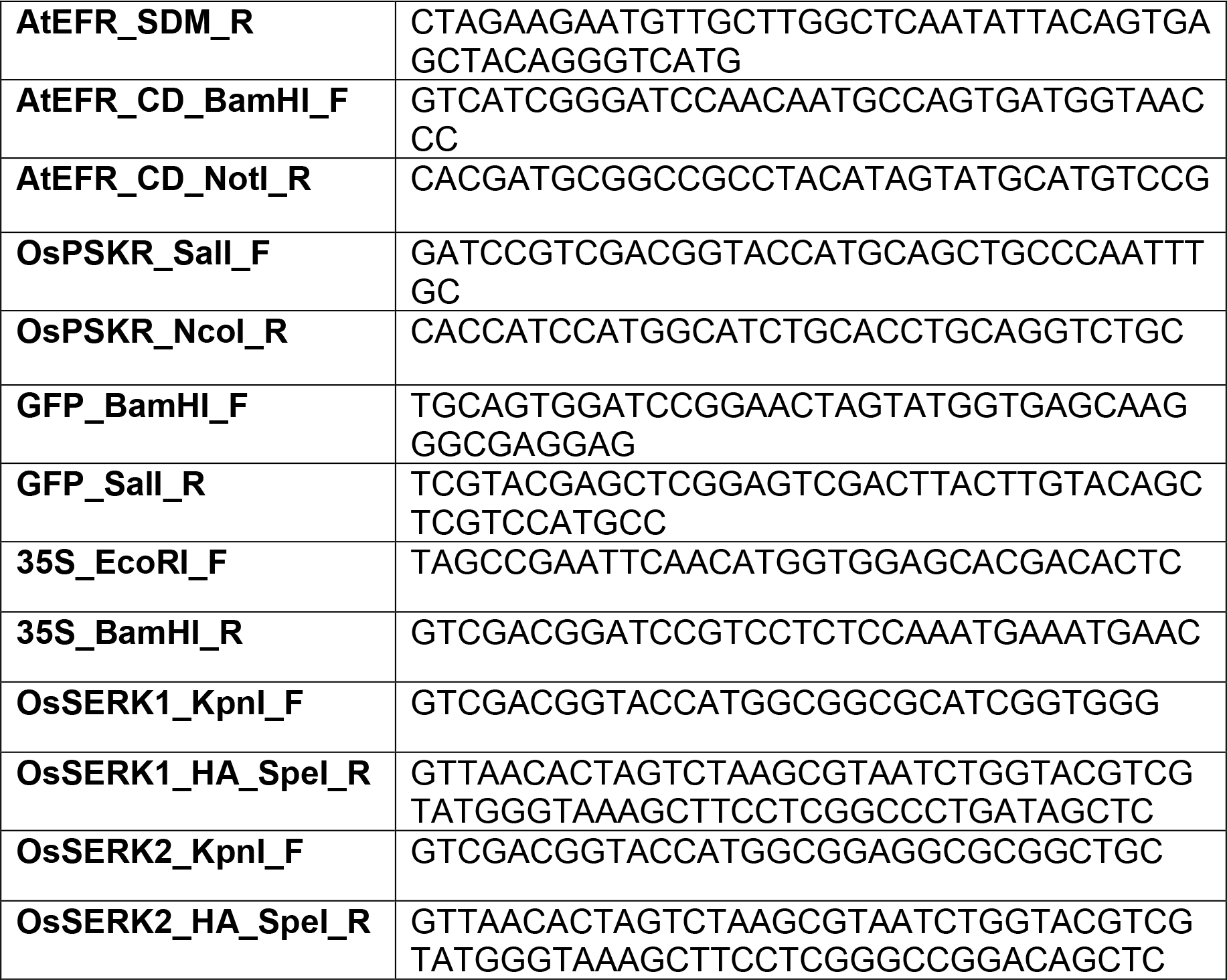
List of primers used in this study.

**Table S2:**
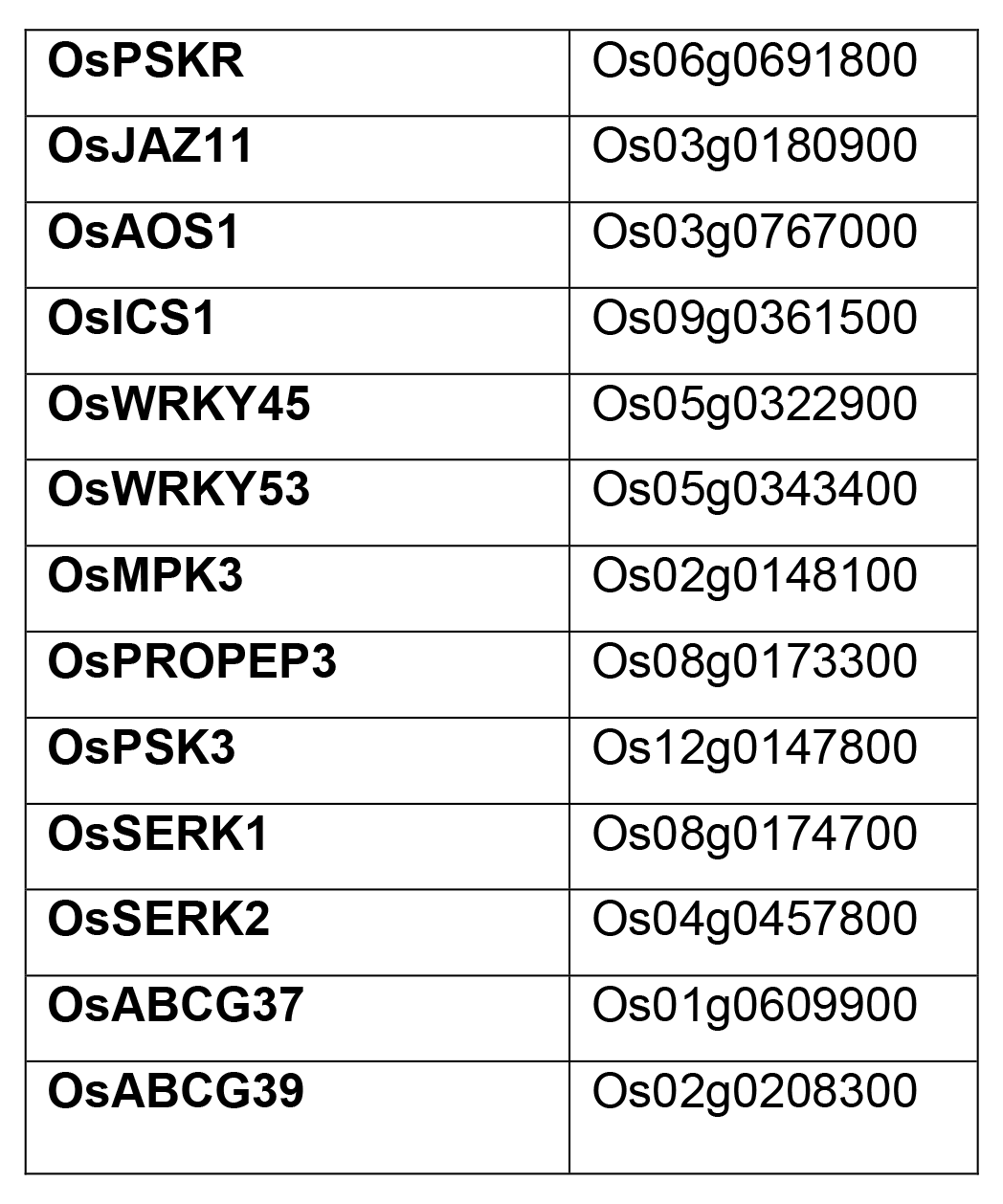
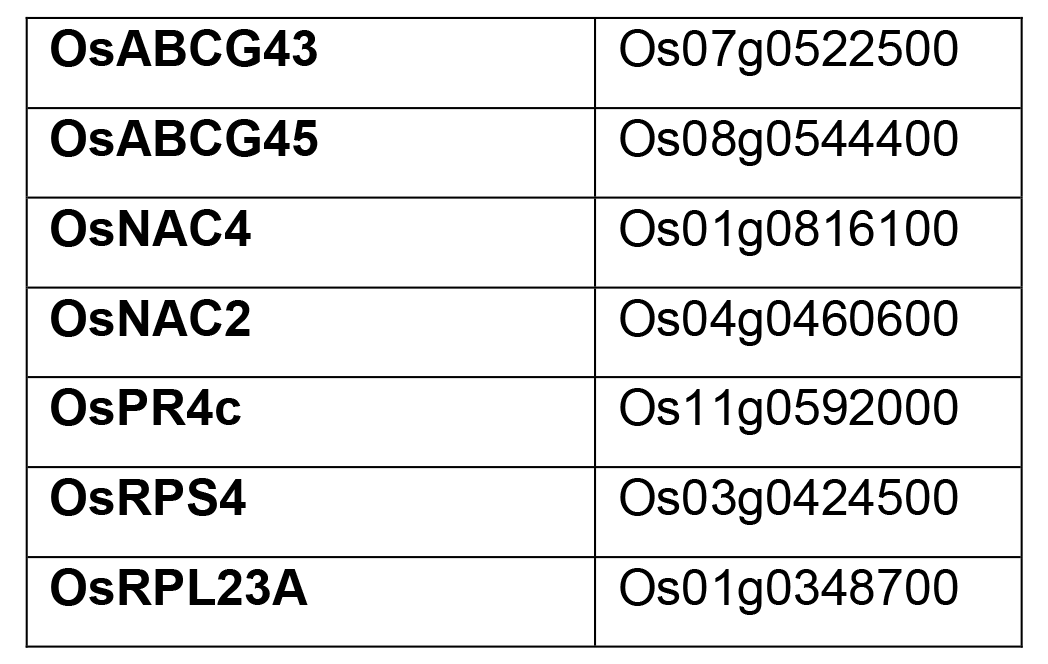
List of Gene IDs of all the genes mentioned in this study.

### Supplemental Datasets

Supplemental dataset 1: List of DEGs with FPKM values upon wounding across different time points in WT plants related to Fig. 1b.

Supplemental dataset 2: Log_2_FC values of the small peptide coding genes upon wounding across time points related to Fig. 1d.

Supplemental dataset 3: List of DEGs with FPKM values upon OsPep2 and OsPep3 treatment across different time points in WT plants related to Fig. 1g.

Supplemental dataset 4: Log_2_FC values of PSK precursors upon wounding as well as OsPep2 and OsPep3 treatment across time points in WT plants related to Fig. 2a.

Supplemental dataset 5: List of DEGs with FPKM values in OsPSKR OE plants, KO plants and KO cell death displaying leaves related to Figs. 5c, 5e and 6c.

Supplemental dataset 6: Log_2_FC values of the 53 wound responsive genes under OsPSKR regulations upon wounding as well as OsPep2 and OsPep3 treatment related to Fig. 5g.

Supplemental dataset 7: Row z-score and FPKM values of DEGs upon wounding at 12 h in WT, OE and KO plants related to Fig. 6e and 6f.

Supplemental dataset 8: Log_2_FC values of predicted PSKRs upon wounding as well as OsPep2 and OsPep3 treatment across time points in WT plants related to Fig. S2a.

